# In-source fragmentation in mass spectrometry-based proteomics: prevalence, impact, and strategies for mitigation

**DOI:** 10.64898/2026.03.27.714398

**Authors:** Thorben Schramm, Ludovic Gillet, Viviane Reber, Natalie de Souza, Matthias Gstaiger, Paola Picotti

## Abstract

Peptide-level analyses are becoming increasingly popular in mass spectrometry-based proteomics and are being applied, for example, in immunopeptidomics, structural proteomics, and analyses of post-translational modifications. In such analyses, peptides that are not biologically meaningful but instead arise as artifacts prior to mass spectrometry analysis pose the risk of data misinterpretation. Here, we describe an approach based on retention time analysis and precise chromatographic peak matching to identify peptides generated by in-source fragmentation (ISF), which occurs between chromatographic separation of peptide mixtures and the first mass filter of a tandem mass spectrometer (MS). To understand the prevalence and properties of ISF, we generated 13 proteomics datasets and analyzed them along with additional 25 previously published datasets spanning a broad range of sample types, MS, and proteomics approaches including classical bottom-up proteomics, immunopeptidomics, structural proteomics, and phosphoproteomics. We found that, in typical trypsin-digested samples on average 1 % of fully-tryptic peptides and 22 % of semi-tryptic peptides originated from ISF. However, we observed large variations between datasets, and in-source fragments exceeded, in some cases, a third of the total peptide identifications. The extent of ISF was dependent on the peptide sequence, the instrument, method parameters, and sample complexity. Although ISF did not impair relative quantification across samples, it generated peptides that could be misinterpreted qualitatively, inflated peptide identifications, and comprised up to 37 percent of peptides shorter than 9 amino acids in immunopeptidomics datasets. We propose that, for peptide-centric applications, our open-source ISF detection approach be used to re-annotate peptides generated by ISF and remove them to avoid misinterpretation of data. ISF is an increasing concern with improving mass spectrometers, as they enable detection of an ever-increasing number of *m/z* features, including low abundance features like ISF products. Our work thus addresses a growing issue in proteomics and presents solutions to mitigate the impact of in-source fragment peptides. In the future, improved feature detection algorithms may enable elucidation of new ISF patterns affecting side chains that have been missed so far, which could contribute to explaining the vast space of as-yet unannotated proteomics data.

## Introduction

Mass spectrometry (MS) has emerged as a standard tool for the analysis of complex metabolite^1,2^ and protein^3,4^ samples. While MS has been broadly applied in scientific and industrial research for decades, an increasing number of MS methods are now applied in clinical diagnostics as well^5^. In MS, analytes such as metabolites and peptides are ionized prior to measurement^6^, with electrospray ionization (ESI)^7^ being one of the most frequently used ionization methods in current approaches. During ESI, a liquid phase containing analytes is sprayed through an injection needle and subjected to high temperatures (25 – 500 °C)^8^ and electric potentials (500 - 4500 V)^6^. This leads to the ionization of the analytes and evaporation of the liquid phase. Although ESI is often described as a *soft* or *gentle* ionization technique^6,9–11^, analytes can be chemically modified or partially fragmented during ESI^11–13^. Such in-source fragmentation (ISF) is typically considered an artifact because it creates new molecular species that can be misinterpreted as biologically relevant.

In the metabolomics field, ISF has been extensively studied^10–18^, and depending on the study, 5 up to 70 % of the data have been reported to originate from ISF^10,11,13,15–18^. Several bioinformatic tools and strategies have been developed to annotate in-source fragments in metabolomics data^10,11,15,19^. If analytes are separated by chromatography before MS measurements, in-source fragments can readily be identified because they share the same retention time and chromatographic elution profiles as their parental molecule^11,19^. In cases where retention times are not available, for example in high-throughput metabolomics approaches that rely on flow injection analyses^20^, network approaches can identify in-source fragments^13,15^.

In contrast to the metabolomics field, ISF has been much less studied in the context of proteomics. A possible reason for this is that the primary goal of classical bottom-up proteomics has been the differential abundance analysis of proteins, which was dominated by data dependent acquisition (DDA) MS methods and by peptide search engines that mostly focus on identifying fully-tryptic peptides. In this context, the probability of detecting fully-tryptic peptides as artifacts arising from ISF is low because for such a mis-annotation to occur, a longer parental peptide arising from missed trypsin cleavage events would have to fragment into a fully-tryptic peptide of sufficient intensity to be selected for MS2 measurements by DDA. In protein abundance analysis, potential effects of ISF on the accuracy of peptide quantification would be diminished during aggregation of peptide quantities to protein quantities. Therefore, ISF has not been considered of major concern so far. In support of this, a previous study identified ISF as a major source of half-tryptic peptides in DDA MS, but showed that it had a low impact for complex samples measured, as in-source fragments represented only 1 - 3 % of the analyzed DDA data^21^. However, data independent acquisition (DIA) MS methods have gained much in popularity in recent years^22,23^. Together with new mass spectrometers with drastically increased sensitivity, they enable the detection of an ever-increasing number of peptides, and it is unclear whether such approaches are more prone to identifying in-source fragments than previous methods. Further, it is known that instrument parameters have an impact on ISF, but the extent of these effects and the relative impact of different parameters is unclear. To complicate this matter, the number of proteomics applications that rely on peptide-centric analyses to quantify half- or non-tryptic peptides, or post-translational modifications (PTMs), is steadily increasing^24^. In these analyses, the quantitative impact of ISF is expected to be more pronounced as peptide quantities would not be aggregated to protein quantities. In addition, qualitative effects of ISF on peptide identification could substantially compromise data interpretation. For instance, in mass spectrometry-based immunopeptidomics or structural proteomics approaches using limited proteolysis^25,26^, biological findings rely on the precise characterization of every peptide from a given protein, including those with semi- or non-tryptic termini. It remains unclear how ISF impacts peptide-centric analyses, quantitatively and qualitatively.

We have therefore revisited the prevalence, characteristics, and impact of ISF in the context of proteomics focusing on DIA datasets acquired with state-of-the-art mass spectrometers. We systematically evaluated a range of instrument parameters and sample types, with particular emphasis on assessing the implications of ISF for peptide-centric proteomics approaches. Further, we provide practical solutions on how to mitigate ISF in proteomics.

## Results

### Detecting in-source fragment peptides in proteomics data

To analyze the prevalence and impact of ISF products, we first sought to develop a bioinformatic pipeline for their detection and quantitative analysis. Peptides are usually separated by liquid chromatography prior to electrospray ionization and mass spectrometry in bottom-up proteomics (Supplementary Fig. 1). Since ISF occurs after chromatographic peptide separation and prior to the first mass filter, in-source fragments and their respective parent peptides have the same retention time profiles despite being of different molecular mass. Fragment peptides also share their amino acid sequence with their parents. We therefore developed an algorithm to detect peptide ISF by identifying peptides with shared sequences that co-elute, or, more accurately for in-source fragments, have the same retention time (RT). To illustrate our approach, we consider a theoretical example with 8 different peptides that share a sequence (Fig 1.a), which yield 28 unique peptide pairs. For each pair, the difference between the apex retention times (ΔRT) was calculated by subtracting the RT of the shorter (or equally sized) peptide (peptide 1) from the RT of the longer peptide (peptide 2) (Fig. 1.b). Peptides that resulted from ISF, which were peptides I, II, VI, and VII in our example, should have small or negligible ΔRTs if paired with their respective ISF parent peptides. We thus identified these peptides by defining a ΔRT cutoff and inspecting which peptide pairs have a ΔRT that fell below it.

**Fig. 1.**
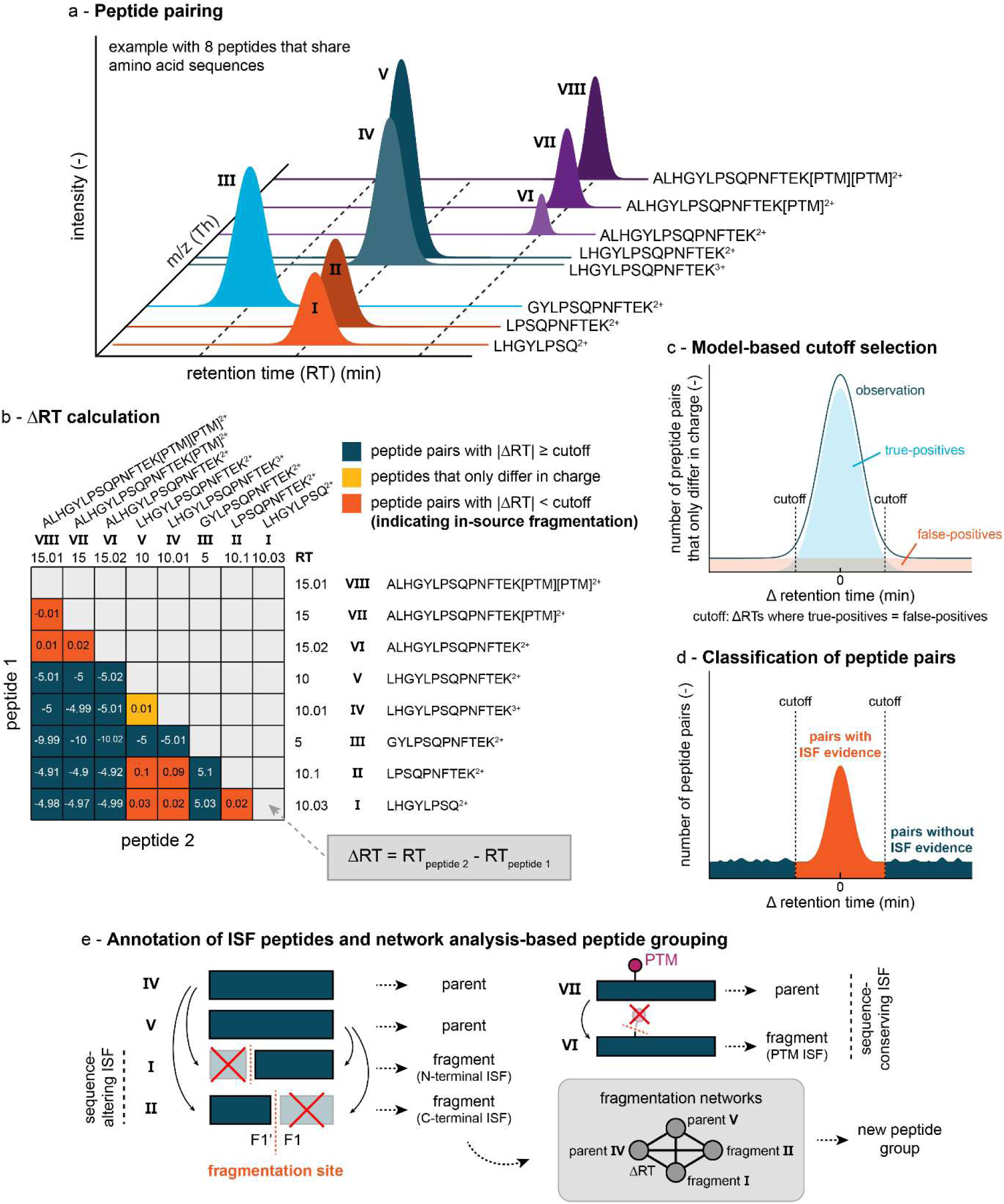
Illustration of the ISF detection algorithm. **a,** Graph showing hypothetical chromatograms of different peptide precursors that share amino acid sequences and thus form peptide pairs. The axes display the retention time, the *m/z*, and the signal intensity as indicated. The peptides form three coeluting groups. **b,** Heatmap illustrating all pairwise combinations of peptides from the example in A and their respective ΔRTs. The ΔRT is calculated by subtracting the apex RT of the shorter peptide from that of the longer peptide. Peptide pairs that do not show co-elution and have ΔRTs larger than a cutoff are blue. Peptide pairs with evidence of ISF are red. Pairs of peptides that only differ in charge are yellow. **c,** Diagram illustrating a theoretical ΔRT distribution of peptides pairs that only differ in charge. An observed ΔRT distribution (blue line) is the sum of two sub-distributions: a normal distribution of true-positive peptide pairs (blue shade) centered around zero and a uniform distribution of false-positive peptide pairs (red shade), of which at least one peptide per pair is falsely annotated. The ΔRT cutoff is the ΔRT at which the value of the normal distribution equals the uniform distribution. **d,** Graph showing a theoretical ΔRT distribution of all peptide pairs that are not part of C. The ΔRT cutoff is used to classify peptide pairs into those with ISF evidence, if they fall within the cutoff, and those without ISF evidence. **e,** Illustration of peptide annotation: peptide pairs with ISF evidence are deconvoluted, and peptide precursors assigned as in-source fragments or parent peptides. Peptides that show ISF often form complex fragmentation networks, in which edges indicate ISF evidence (labelled with the respective ΔRT) and nodes are peptides. Fragmentation networks can be used to create new peptide groups. Amino acid sequence-altering ISF can occur N-terminally, C-terminally, or at both ends (omitted in the illustration). ISF of exclusively PTM groups are examples of amino acid sequence-conserving ISF.

As RT distributions, chromatographic peak shapes, and resolution can vary between instrumental setups or runs, we determined the “co-elution” ΔRT cutoff for each sample individually. To obtain this cutoff, we analyzed the data at the peptide precursors level, as throughout this study if not stated otherwise, and used the ΔRT distribution of peptides that differ only in charge and not in mass, which were peptides IV and V in our example. Because these peptides must have the same RT, they provide ground truth for the natural variation of ΔRT values of co-eluting peptides (Fig. 1.c). As false-discovery-rates are commonly set at 1 % for peptide searches, a small fraction of peptide pairs will contain a falsely annotated peptide resulting in a continuous uniform distribution of ΔRT values corresponding to these false positives. The ΔRT cutoff for co-eluting peptides is defined as the ΔRT at which the normal distribution of true-positives intersects with the continuous uniform distribution of the false-positives. In the next step, peptide pairs were classified as pairs with ISF evidence, if their ΔRT was below the ΔRT cutoff, or as pairs without ISF evidence (Fig. 1.d). In the last step of our ISF detection, peptide pairs were uncoupled and, in case of ISF evidence, shorter, type 1 peptides annotated as fragment peptides, and longer, type 2 peptides as parent peptides (Fig. 1.e). If there was no ISF evidence, peptides were referred to as non-ISF peptides.

In case of a parent peptide yielding more than one fragment peptide, the intermediary fragments will still be annotated as fragments but not as parent peptides, implying that parent and fragment peptide annotations are mutually exclusive. Multiple in-source fragments of a parent peptide also enable the construction of ISF networks, in which each peptide is a node and each edge a peptide pairing with a ΔRT. These fragmentation networks enable the identification of in-source fragments that share the same parent and can be used to define new peptide groups, for which all individual peptide quantities of each group can be aggregated to single peptide group quantities. We further identify C-terminal ISF if the N-terminal part of the parent is detected as a fragment, and *vice versa* for N-terminal ISF. We denote the first amino acid at the C-terminal side of the ISF site F1, and the first amino acid at the N-terminal side F1’, analogous to the nomenclature of proteolytic cleavage sites (P1, P1’, etc.). While the detection of F1 and F1’ implies an ISF that alters the amino acid sequence of the parent peptide, it is in principle possible that an ISF does not change the amino acid sequence but affects a side chain or PTM. As some PTMs like oxidation of methionine are included in most peptide searches by default, we also consider PTM in-source fragmentations (PTM ISF), defined as an ISF event that detaches the PTM without altering the amino acid sequence.

We tested our ISF detection algorithm on DIA data from a *Saccharomyces cerevisiae* 288c sample spiked with 8 proteins, which we used as a reference sample throughout this study. We chose this sample as a reference because we suspected that peptides with high concentrations were more likely to yield measurable in-source fragments, and this sample yielded such peptides (from the spiked in proteins) in a controlled fashion without compromising on overall sample complexity. Our model identified a ΔRT cutoff of ± 0.129 min for co-eluting peptides with a recall of 97.1 % for the true positive peptide pairs, which were the peptides of equal mass but different charge states (Supplementary Fig. 2a). This ΔRT cutoff was more conservative than the median full width at half maximum (FWHM), which was 0.19 min and is often used to determine peak separation. Next, we used the ΔRT cutoff to determine ISF peptides and found that 24.4 % of all peptide pairs that differed in molecular mass fell within the cutoff (Supplementary Fig. 2b). This result provides the first evidence that in-source fragmentation is prevalent in DIA proteomics data.

In the ΔRT distribution, we further observed a broad peak of peptide pairs around -10 min. We found that this was due to peptides carrying an oxidation group as PTM (Fisher’s exact test checking for enrichment between -13 and -5 min, right-tailed, *P*-value < 1e-200). We also observed a long tail of the ΔRT distribution for ΔRT > 0, which was expected because peptide pairs that were not due to ISF typically have a longer type 2 peptide that is often stronger retained on the chromatographic column than the shorter type 1 peptide.

To our knowledge, there are no published, open access software tools to detect ISF in DIA proteomics data to which we could compare our ISF detection algorithm. However, we compared it to another, commercially available and unpublished algorithm that was developed in parallel to our study as a feature for Spectronaut 20 (Biognosys). For our yeast reference sample, 97.7 % of all ISF annotations across n = 4 technical replicates (4716 out of 4828) were identical, showing broad agreement between both algorithms (Supplementary Fig. 3) and providing a cross-validation.

### In-source fragmentation is abundant

Next, to determine how abundant peptide ISF is in data from typical trypsin-digested bottom-up proteomics samples measured by DIA, we analyzed 16 published datasets, acquired with 9 different mass spectrometers models, from different research groups (Supplementary Tab. 1)^27–42^. These datasets comprised 317 samples from a broad range of sources, including *Escherichia coli*, *Saccharomyces cerevisiae*, *Arabidopsis thaliana*, *Mus musculus*, and *Homo sapiens,* and comprising both laboratory cultures and human biopsies. The number of detected protein groups ranged from 3 to 6,180. In-source fragments made up 3.4 ± 3.2 % of the total peptide precursor identifications (mean and standard deviation, n = 317 samples) across the datasets, and 4.8 ± 4.2 % of the total peptide intensity (Fig. 1a). However, we observed a large variation in the extent of ISF, ranging from 0.2 ± 0.1 % by intensity (PXD062917, n = 9 samples; 0.5 ± 0.5 % by identifications) to 15.5 ± 0.3 % by intensity (PXD069457, n = 6; 13.1 ± 0.1 % by identifications).

Increasing sample complexity generally yielded lower shares of in-source fragments within a dataset (% by identifications and by intensities), although not every sample with a low complexity had high levels of ISF. To better understand how sample complexity affects ISF, we prepared the following 7 trypsin-digested samples of ascending complexity and compared the prevalence of ISFs: (1) mix of 11 peptides, (2) mix of 8 proteins, (3) *E. coli* BW25113 lysate, (4) affinity-purification MS (APMS) from a *H. sapiens* kinase pull-down (5) *S. cerevisiae* S288c lysate, (6) *H. sapiens* HEK-293 lysate, and (7) the tribrid proteome sample obtained by combining samples 3, 5, and 6. We detected ISF peptides in all samples (Fig. 2.b). The tribrid proteome sample (n = 4 technical replicates) had the lowest level of ISF, with fragments representing 4.01 ± 0.06 % of total peptide intensity (2.591 ± 0.005 % of identifications, mean and standard deviation), while the mix of 8 proteins had the highest value (24.1 ± 1.3 % by intensity; 31.0 ± 0.5 % by identifications). Across all samples, both the intensities as well as the numbers of fragment peptide identifications increased with decreasing sample complexity. The absolute number of parent peptides was on par with the number of fragments, except for the low complexity samples (peptide and protein mixes), which had more fragments than parents (Fig. 2.c). Although we observed fewer parent peptides than non-ISF peptides, the intensities of the parent peptides were in the same range as the non-ISF peptides (Fig. 2.c). This also meant that at least 32 % (*H. sapiens*) and up to 95 % (mix of peptides) of the total peptide intensity could be associated with ISF, combining the intensities of fragments and parents (Fig. 2.d). This was consistent with our observations in the published datasets, in which on average 37 ± 22 % of the total peptide intensity could be associated with ISF, with large variation between individual datasets. Across all datasets (published and our new data reflecting a range of complexity), 82 % of the in-source fragments were semi-tryptic peptides (Fig. 2.e), and in-source fragments represented 0.2 – 83.3 % of semi-tryptic peptides, depending on the sample (Fig. 2.f).

**Fig. 2.**
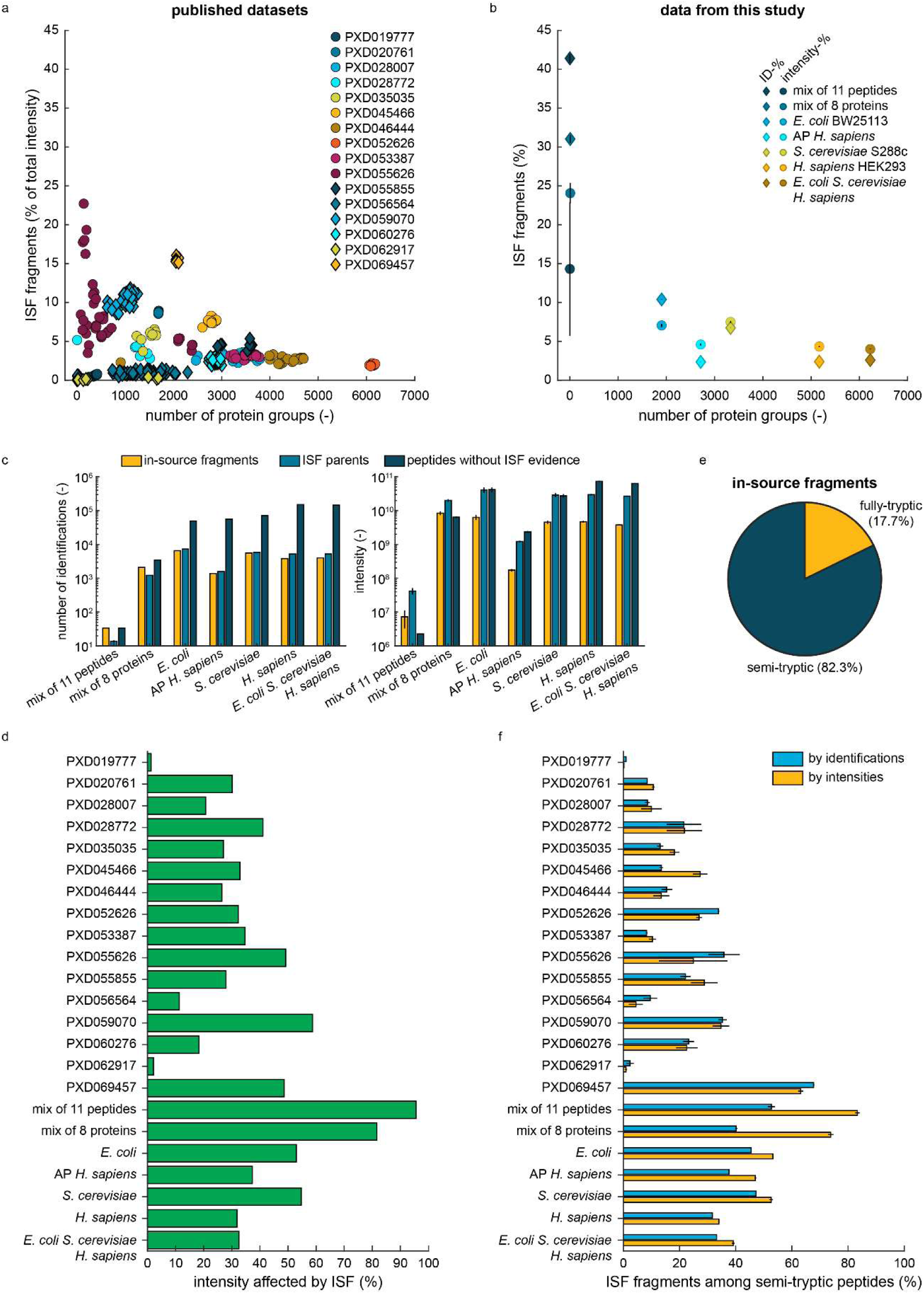
Evidence for abundant ISF across proteomics data of different complexity. **a,** Dot plot showing the share of in-source fragments (% by intensity) per number of detected protein groups (minimum of 5 unique peptide sequences per group) in 317 samples across 16 published proteomics datasets (as indicated by color and symbol shape). **b,** Dot plot showing the share of in-source fragments (dots: % by intensity, squares: % by identifications) per number of identified protein groups (minimum of 5 unique peptide sequences per group) across 7 datasets (as indicated by colors) that ranged from a mix of 11 peptides up to a mix of three whole proteomes. Dots and squares are mean values (n = 4), vertical lines indicate standard deviations. **c,** Bar graphs showing the number of identifications and the absolute intensity of in-source fragments, ISF parents, and peptides without ISF evidence for the datasets shown in B. Bars indicate mean values (n = 4), vertical lines standard deviations. **d,** Bar plot showing the percentage of total intensity covered by in-source fragments and parents combined. **e,** Pie chart showing the share (%) of semi- and fully-tryptic peptides among in-source fragments across all the datasets from A. **f,** Bar plot depicting the average share of semi-tryptic in-source fragments (%) among all detected semi-tryptic peptides per dataset from A. Blue bars are intensity-based representations (yellow bars: number of identifications-based). Bars indicate mean values (n = 4), horizontal lines the standard deviation.

Taken together, these results showed that ISF varies strongly between datasets, depends on overall sample complexity, and can account for a large share of the mass spectrometry signal. ISF is especially prevalent in low complexity samples, highlighting the importance of ISF detection in those cases.

### Characterization of ISF peptides

To learn general properties of the observed ISF peptides, we analyzed combined data with 2,407,805 identified peptides from all the (published and newly generated) datasets described above. 30.3 % of the peptides shared their sequence with another peptide and formed around 13.7 million peptide pairs (Fig. 3.a). 1,222,589 of the peptide pairs (8.9 %) showed evidence of ISF, of which 67.9 % had sequence-altering ISF, and 26.6 % were fragment-to-fragment pairs. Most fragment peptides originated from N-terminal ISF of fully-tryptic peptides and had a C-terminal arginine or lysine (Supplementary Fig. 4.a). Further, non-ISF peptides had mostly lower intensities than in-source fragments and parents, with parent peptides having higher intensities than fragments (Fig. 3.b and Supplementary Fig. 4.b). This indicated that the detected in-source fragments and parents either derived from abundant proteins, were especially amenable to measurement by MS, or both. As expected, in-source fragment peptides often had a lower number of charges than their respective parent peptides and were shorter than the parent and non-ISF peptides, as 47.3 % of the ISF peptide pairs showed losses in charge and 92.5 % showed losses of amino acids due to ISF (Supplementary Fig. 4.c, d, e, and f).

**Fig. 3.**
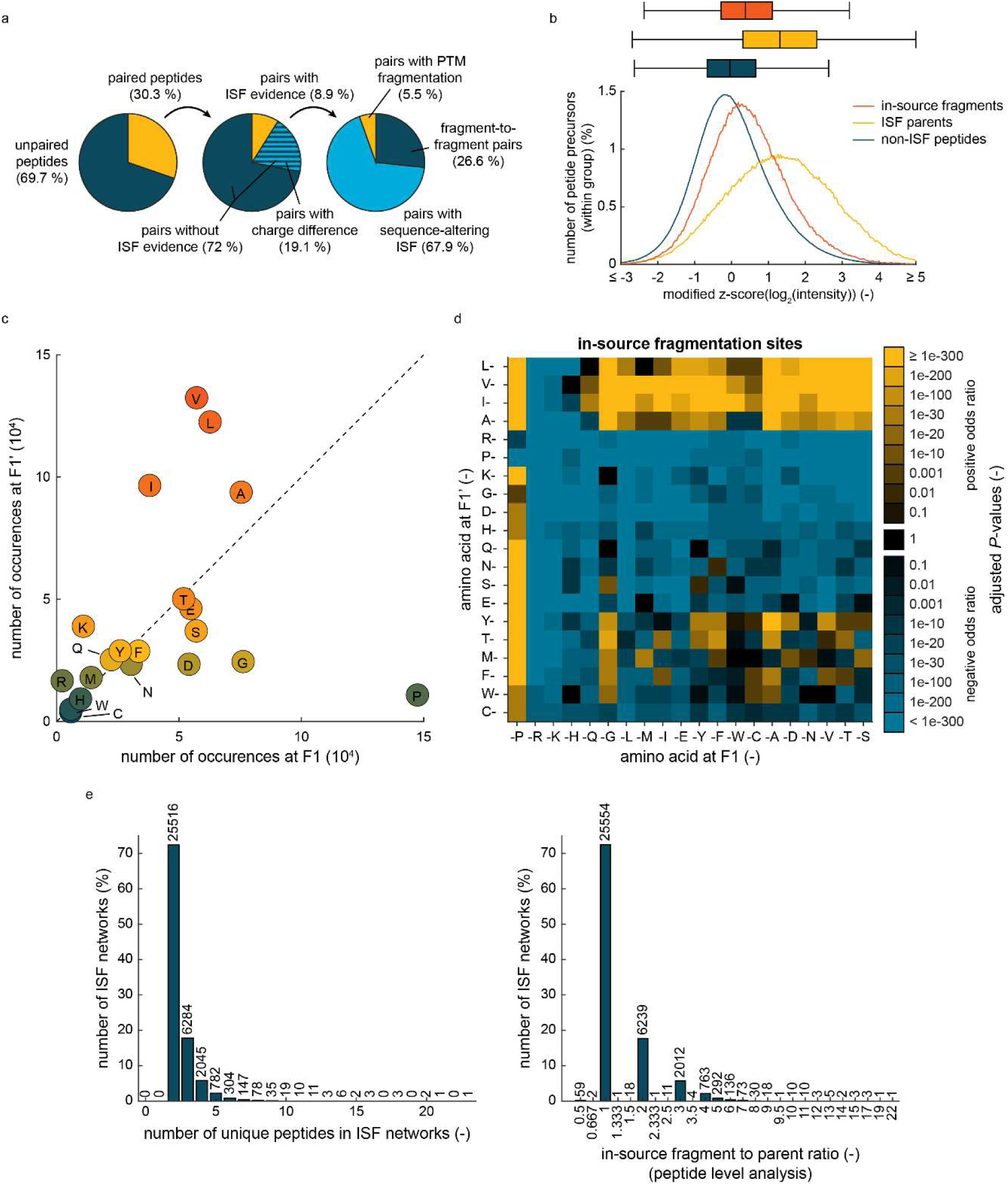
ISF peptide characterization. **a,** Three pie charts illustrating: (1) the fractions of peptides with and without peptide pairing (in %), (2) the number of peptide pairs with and without evidence of ISF (%), and (3) the number of peptide pairs with ISF evidence that have a sequence-altering ISF, are fragment-to-fragment pairs, and feature PTM ISF (%). The data used for this figure stems from 23 datasets (also see Fig. 2) that were merged. **b,** Graph showing the z-scored (modified) log_2_-intensity distributions (in %) of in-source fragment (red) and parent (yellow) peptides as well as peptides without ISF evidence (blue). Box whisker plots above the distributions indicate the median values and interquartile ranges. Modified z-scores are calculated with median values and median absolute deviations. **c,** Dot plot showing the abundance of amino acids at the F1’ position at ISF sites over occurrences at F1. **d,** Heatmap showing the adjusted *P*-values from an enrichment analysis (Fisher’s exact test, two-tailed, *P*-value adjustment by Benjamini-Hochberg procedure^43^) of all peptide bonds observed at IFS sites. Blue squares indicate a negative odds ratio, yellow a positive ratio. **e,** Histograms showing the number of ISF networks (in %) with the indicated number of unique peptides per network (left) or with the indicated in-source fragment to parent ratio (right).

To understand if ISF events had a bias for certain amino acid bonds, we analyzed how often an amino acid occurred at F1’ or F1 of an ISF site. Valine followed by leucine, isoleucine, and alanine were most abundant at F1’, while proline was most abundant at F1, followed by glycine and alanine (Fig. 3.c). An enrichment analysis of peptides bonds at ISF sites, using all possible peptide bonds of detected peptides as null distribution (two-tailed Fisher’s exact test), showed that 56 out of 80 possible peptide bonds with short, hydrophobic amino acids (valine, isoleucine, leucine, and alanine) at F1’ showed a significant enrichment (Benjamini-Hochberg^43^ adjusted P-values < e-20), and 17 out of 20 bonds with proline at F1 were enriched at ISF sites (Fig. 3.d). The aspartate-proline bond is known to be susceptible to non-enzymatic cleavage under acidic conditions^44^. Here, we did not observe an enrichment of the aspartate-proline bond at fragmentation sites but rather a depletion. However, we detected enrichment of the aspartate-proline bond at non-tryptic termini of semi-tryptic non-ISF peptides (Supplementary Fig. 5).

As they made up over a quarter of the ISF peptide pairs, fragment-to-fragment pairs indicated that multiple ISF events of peptides are abundant. We thus constructed ISF networks with sequence-altering ISFs to better understand this effect and its prevalence but also to assign ISF peptides to new peptide groups. Merging information from peptides with multiple charges, we observed that 72.4 % of the ISF networks had two unique peptide sequences and 72.5 % a one-to-one ratio between in-source fragment and parent peptides (Fig. 3.e), implying that the majority of ISFs were from a single ISF parent to a single fragment peptide. However, the remaining 27.6 % of the ISF networks contained more than two unique peptide sequences, with the largest ISF network having 23 unique peptide sequences (Supplementary Fig. 6). 27.3 % of the ISF networks also had an imbalance between the number of in-source fragments and parent peptides (Fig. 3.e). Detecting both N- and C-terminal fragments from a single ISF event appeared to be challenging as we only observed 9 cases across the 23 datasets, in which the combined sequences of two in-source fragments matched the parent sequence.

These results show that peptide ISF mostly occurs at specific peptide bonds prone to fragmenting in the gas phase and often creates a complex pattern of peptide products and *m/z* features, which can be deciphered by network analysis.

### In-source fragmentation of PTMs

If ISF only removes the PTM group of a peptide, the resulting fragment peptide can appear as an unmodified peptide, which could be mistaken for an authentic identification of the unmodified peptide. We analyzed how often this effect occurs for several PTMs.

Methionine oxidation and N-terminal acetylation are commonly observed PTMs in proteomics data; we included them as variable modifications in the peptide searches of our combined dataset and studied if they undergo ISF. We detected that ISF of these PTMs comprised 5.5 % of the total ISF events (Fig. 3.a) and that the intensity differences between in-source fragments and parents were much smaller for PTM ISF than for sequence-altering ISF (Supplementary Fig. 4.b). Most of the cases involved methionine oxidation, and only a few N-terminal acetylation (0.4 % of all PTM ISF) (Fig. 4.a). While there was a large variation in the number of methionine oxidation ISF between the datasets, we did not observe a correlation with the number of peptides carrying a methionine oxidation (Fig. 4.b). This indicated that the amount of PTM ISF was dependent on other factors, e.g. instrument parameters.

**Fig. 4.**
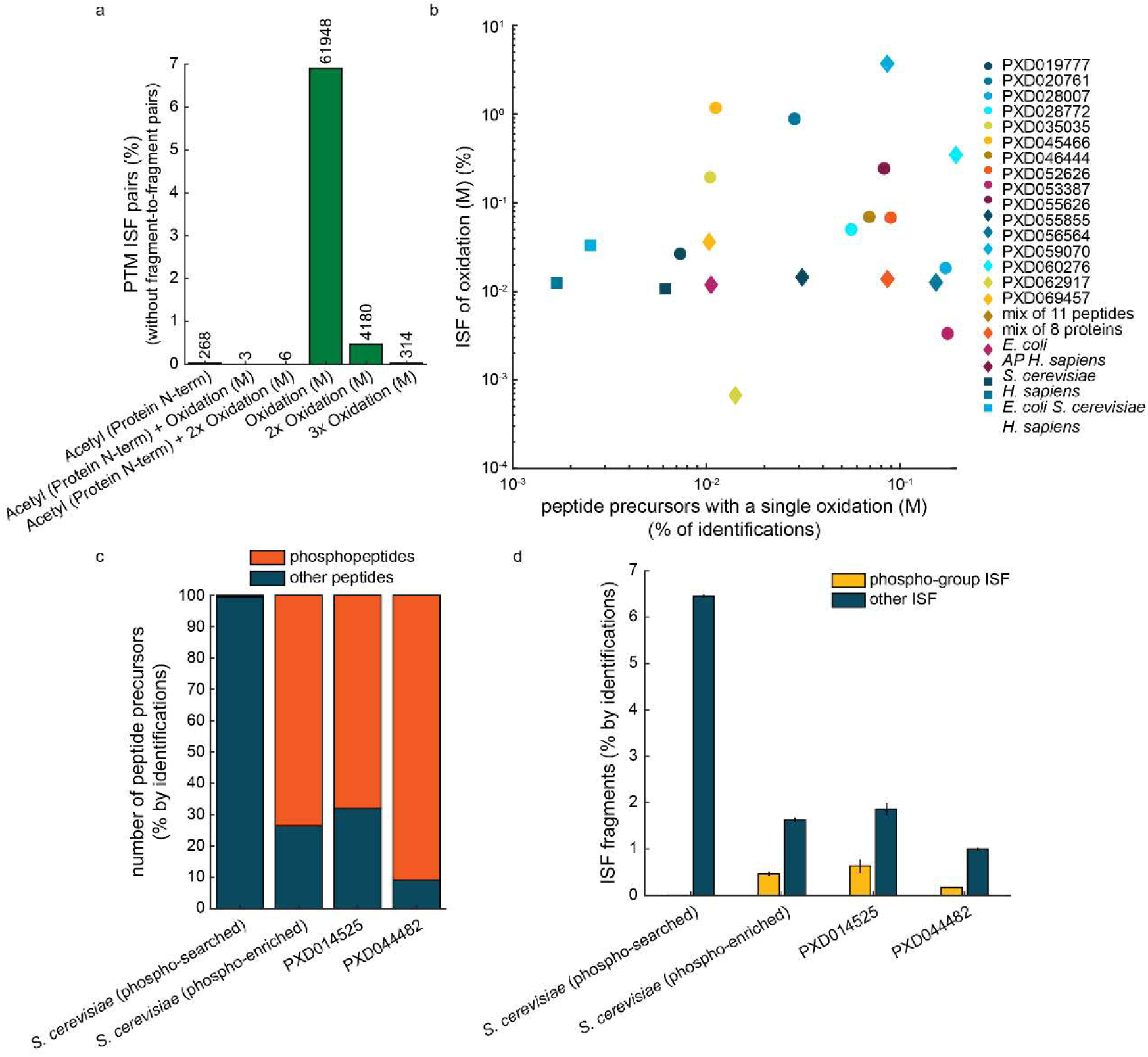
PTM in-source fragmentation. **a,** Bar plot showing the number of peptide pairs with a PTM ISF (as labelled) among all pairs with ISF evidence from 23 combined datasets (also see Fig. 3.). **b,** Dot plot showing the number of ISF of methionine oxidation (%) per number of precursors with a single methionine oxidation (%). Colors and symbols indicate 23 different datasets. **c,** Bar plot showing the percentages of phosphopeptides (red) and other peptides (blue) in four different datasets. The two *S. cerevisiae* datasets were acquired from the same sample source: one as a standard yeast proteomics dataset and the other as phosphoproteomics dataset (including phosphopeptide enrichment). The two literature datasets (PXD014525 and PXD044482) relied on phosphopeptide enrichment. **d,** Bar graph displaying the number of in-source fragments (% by identifications) in the four datasets of C.

It is known that phosphorylated metabolites like ATP tend to lose phospho-groups due to in-source fragmentation^11^, and we thus assessed the prevalence of ISF events affecting phosphopeptides. We first directly re-searched our *S. cerevisiae* S288c dataset for phosphopeptides. Second, we analyzed *S. cerevisiae* S288c digests after phosphopeptide enrichment by titanium dioxide chromatography^45^, and third, we included two additional, published datasets that used various phosphopeptide enrichment strategies (PXD014525^46^ and PXD044482^47^). As expected, the number of phosphopeptides were much higher in the three phospho-enriched datasets, where they made up 68 - 91 % of all identified peptides as compared to 0.6 % in the standard proteomics dataset (Fig. 4.c). Similarly, the number of ISF events leading to loss of a phospho-group was higher in the phosphoproteomics datasets (0.17 - 0.63 % of in-source fragments by identifications, 0.46 – 1.06 % by intensity, mean values, number of samples n = 4 (phospho-enriched yeast dataset), n = 18 (PXD014525), and n = 10 (PXD044482)) than in the standard proteomics dataset (0.0048 ± 4e-5 % by identifications, 0.0037 ± 1e-4 % by intensity, mean and standard deviation, n = 4 technical replicates) (Fig. 4.d). In the two published datasets, we observed a bias of phospho-group ISFs towards phosphorylated tyrosines (Y) indicating that this amino acid is more prone to phospho-group ISF than phosphorylated serines (S) and threonines (T) (Supplementary Fig. 7).

In conclusion, PTM ISF occurs at a detectable level in proteomics data and varies in a dataset and PTM-dependent manner, although it was less prevalent than ISF of regular peptide bonds.

### Peptide ISF is instrument and parameter dependent

The ion funnel design and its radiofrequency (RF) are known to have an impact on the ISF of peptides^48^, the temperature of the ion transfer capillary and the RF affect lipid ISF^49^, and increasing accumulation times during trapped ion mobility spectrometry (TIMS) time-of-flight mass spectrometry (TOF) also increases ISF^50^. Based on this information, we expected ISF to vary between different mass spectrometers. To assess the extent of such variation, we measured a trypsin-digested yeast reference sample mixed with 8 pure proteins on three Thermo Scientific mass spectrometers, an Orbitrap Exploris 480, an Orbitrap Astral, and a Q Exactive Plus Hybrid Quadrupole-Orbitrap (QE+), each using LC and DIA methods based on previously established workflows^51,52^. With 1.07 ± 0.02 and 1.51 ± 0.02 % (mean and standard deviation, n = 4 technical replicates), the QE+ and the Astral mass spectrometers yielded a much lower fraction of peptide identifications corresponding to in-source fragments than the Exploris 480 (5.68 ± 0.03 %) (Supplementary Fig. 8). This was also reflected in the total intensity of in-source fragments (QE+: 2.2 ± 0.3 %, Exploris 480: 8.8 ± 0.2 %, Astral: 3.70 ± 0.05 %). Thus, ISF was indeed highly dependent on the instrument.

As the RF and the ITC temperature were shown to affect ISF^48,49^, we subsequently quantified the impact of these two parameters as well as the electrospray voltage on peptide ISF using the Orbitrap Exploris 480. We used the yeast reference sample again and measured it at a spray voltage fixed at 2500 V with seven variations of the funnel radio frequency and six variations of the ITC temperature. We also measured the same sample with a relative funnel radio frequency fixed at 50 % with six variations each of the other two parameters (ITC temperature and spray voltage), which amounted to a total of 72 unique sets of parameters tested.

The chosen parameters resulted in a broad range of ISF, from low (0.2 % of peptide precursor identifications across all samples, 301 fragments at 100 °C, 2500 V, 10 % RF, n = 1 replicate) to high (4.9 %, 6566 fragments at 350 °C, 2500 V, 60 % RF) (Fig. 5.a and Supplementary Fig. 9.a). Increasing the funnel RF increased both the relative intensity and the number of in-source fragments (Fig. 5.b). Simultaneously, it only increased the overall signal up to a RF of 40 %, indicating that higher funnel RF values are disadvantageous. The effect of the ITC temperature on ISF was dependent on the funnel RF and led to increases in ISF (in signal and identifications) at high RF values but had little effect on ISF at low RF values. In contrast to both other parameters, the electrospray voltage had little effect on overall signals, number of peptide identifications, and ISF, except for the very low setting of 1500 V that decreased overall signal drastically.

**Fig. 5.**
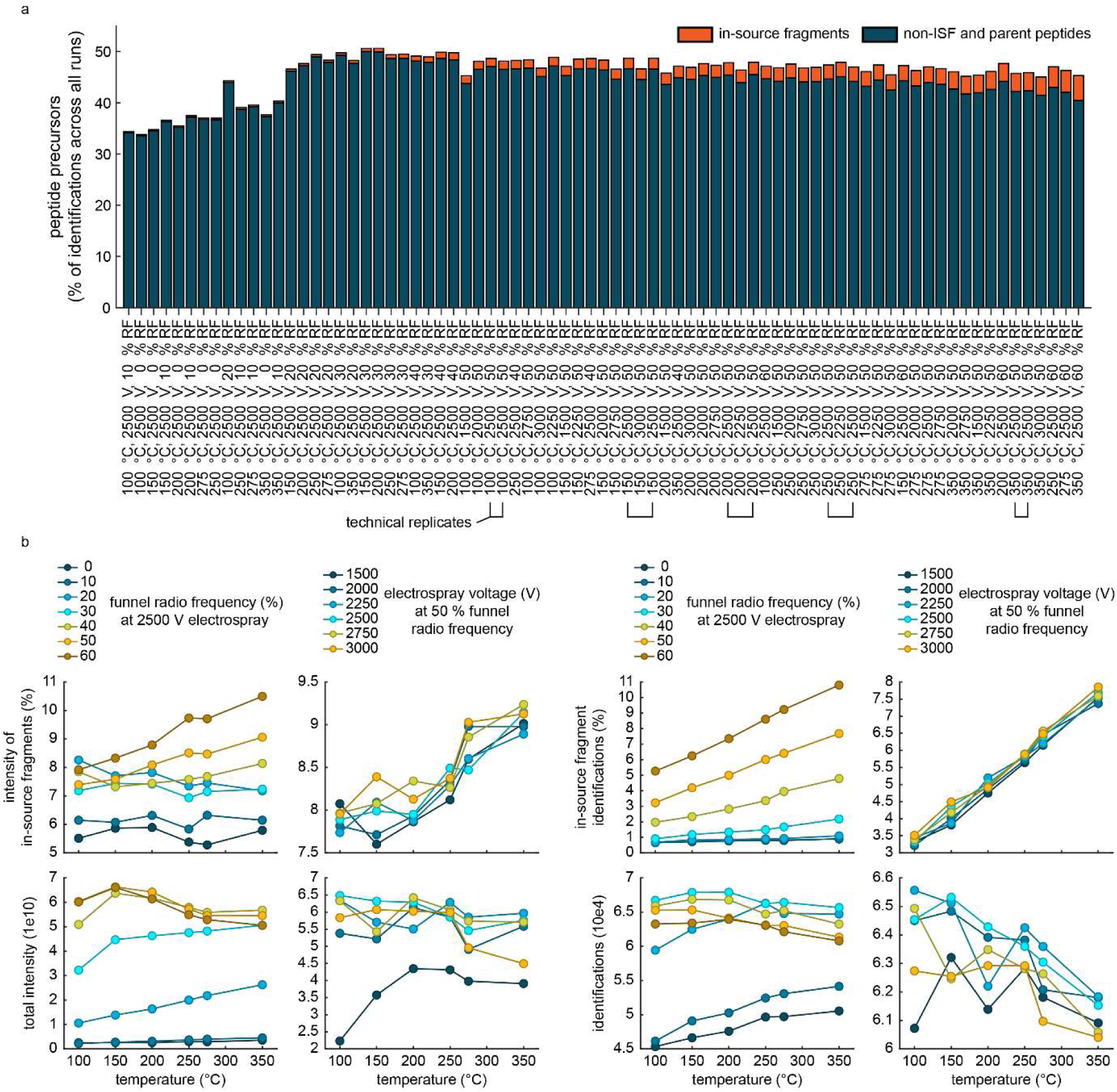
Impact of the funnel RF, electrospray voltage, and ITC temperature on peptide ISF. **a,** Bar plot displaying the number of detected peptide precursors that are non-ISF and parent peptides (blue bars) and that are in-source fragments (red bars) (% of all identifications across all samples) in a yeast reference sample acquired at different funnel radiofrequencies (%), electrospray voltages (V), and ITC temperatures (°C) (n = 1, except 5 pairs of technical replicates as indicated). The samples are sorted from left to right in ascending order of increasing ISF. **b,** Dot plots showing the relative intensity (%) (upper left ), the total intensity (lower left), the relative number of identifications (%) (upper right ), and absolute number of identifications (lower right ) of in-source fragments in the samples from A.

The number of precursor identifications is often used as a quality control measure during data acquisition. Because instrument parameters increase the number of in-source fragments while decreasing the number of identifications of genuine peptides with no evidence of fragmentation, not accounting for ISF will lead to an overestimation of identifications and can thus lead to choosing suboptimal conditions (Fig. 5.a).

In conclusion, these results show that ISF was strongly dependent on the instrument, funnel RF, and ITC temperature, indicating that proteomics experiments can be optimized to avoid high ISF rates. Since ISF can lead to an overestimation of the total peptide identifications, minimizing it should generally be part of method optimization efforts alongside the optimization of other parameters like total peptide identifications and signal intensities.

### ISF does not impair relative quantification

To test if ISF has an impact on relative quantification of peptide precursors, peptide groups, and protein groups, we conducted an experiment in which we mixed, at equal volumes, a yeast sample at a fixed concentration with 8 samples each with a different concentration of *E. coli* peptides (Fig. 6.a). Peptide precursor fold changes were calculated relative to the sample with the highest *E. coli* proteome concentration.

**Fig. 6.**
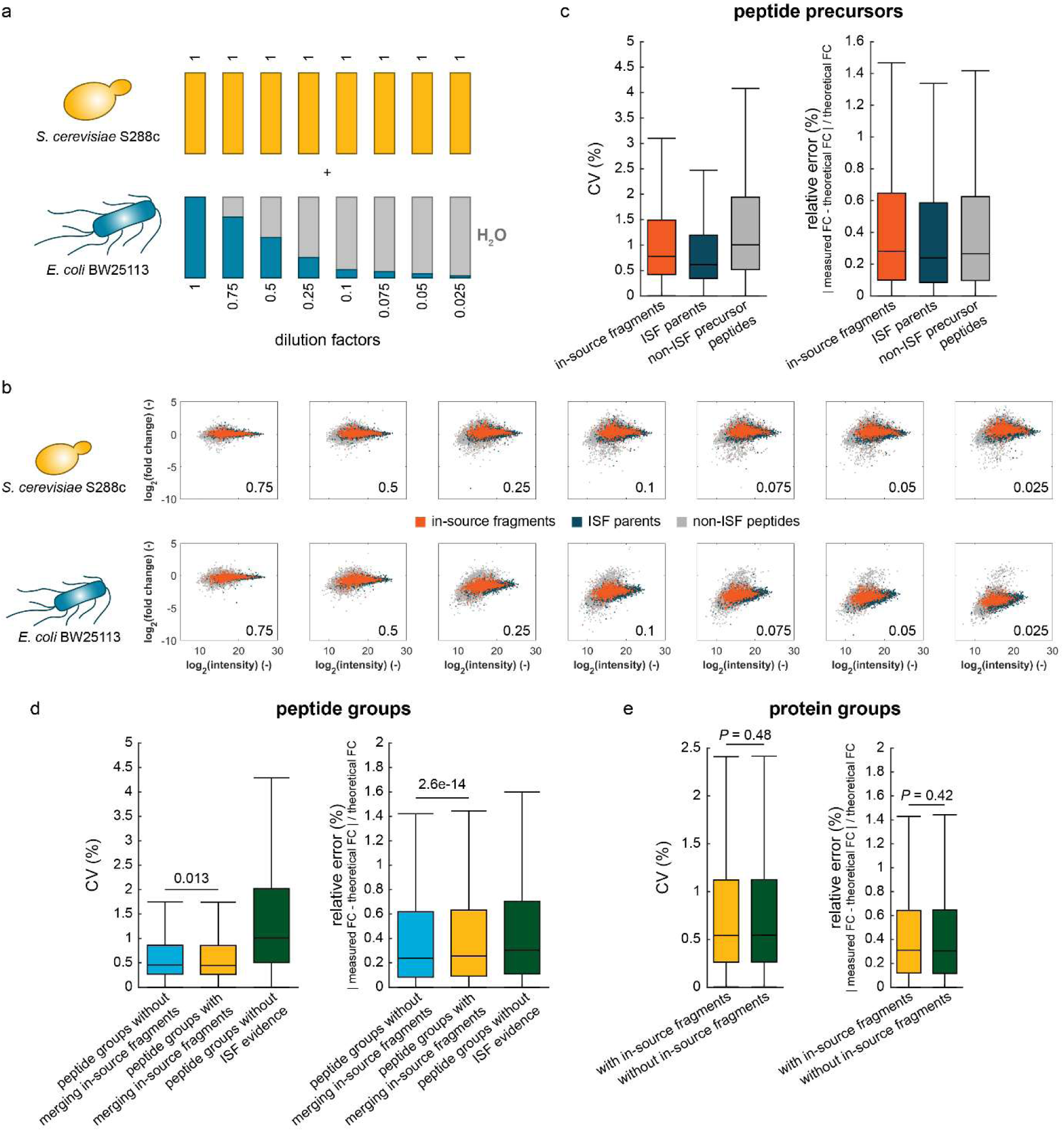
Impact of ISF on quantification. **a,** Whole proteome digests of *S. cerevisiae* S288c and *E. coli* BW25113 were mixed at different ratios as indicated as dilution factors below the blue bars. The yeast proteome concentration was kept constant. The *E. coli* proteome concentration was varied, and volumes compensated with water. **b,** Dot plots showing the mean log_2_-precursor fold changes (relative to the sample with the undiluted *E. coli* proteome) per mean log_2_-precursor intensity (n = 3 technical replicates). Red dots indicate in-source fragments, blue dots ISF parents, and grey dots non-ISF peptides. **c,** Box whisker plot showing the median coefficient of variation (CV, %, n = 3 technical replicates) and the median relative error between theoretical and measured fold changes (n = 3 technical replicates) of all peptide precursors from the samples in B. Data of in-source fragments is indicated in red (parents: blue, non-ISF precursors: grey). Boxes depict interquartile ranges, and whiskers extend to 1.5-fold of the interquartile ranges. **d,** Box whisker plot showing the median coefficient of variation (CV, %, n = 3 technical replicates) and the median relative error between theoretical and measured fold changes (n = 3 technical replicates) of all peptide groups from the samples in B. Peptide groups of all ISF parents without (blue) and with (yellow) merging with their respective in-source fragments are compared with all peptide groups without ISF evidence (green). Boxes depict interquartile ranges, and whiskers extend to 1.5-fold of the interquartile ranges. *P*-values were calculated using the two-tailed Wilcoxon rank sum test. Peptide quantities were calculated by summing up precursor intensities. **e**, Same as D but depicting protein groups that contain (yellow) or omit (green) in-source fragments. Protein quantities were calculated by summing up precursor intensities.

For all samples, the yeast proteome remained stable around a fold change of 1, whereas the *E. coli* peptide precursors followed the dilutions (Fig. 6.b). At lower concentrations of the *E. coli* proteome, the peptide precursor fold changes did not match the theoretical values as accurately as at higher concentrations, which was probably due to ion suppression and poor peptide detectability. However, comparing the data between non-ISF, fragment, and parent peptide precursors revealed that the non-ISF peptide precursors had broader fold change distributions than the other peptide precursors (Fig. 6.b and Supplementary Fig. 10) and that coefficients of variation (n = 3 technical replicates) of non-ISF peptide precursors were often larger than those of in-source fragments and parents (Fig. 6.c, left plot, *P* < e-200 for all pairwise comparisons, Wilcoxon rank sum test, two-tailed). Because the dilution factors were known, we could calculate a relative error between the measured fold changes of the precursor peptide quantities and the expected, theoretical fold changes. These relative errors showed that the non-ISF precursor peptides recovered the expected, theoretical values better than the in-source fragments by only a small margin (0.015 difference in the median relative error); interestingly, the ISF parents recovered the theoretical values the best, which is probably due to their generally good measurability and high signals (Fig. 6.c, right plot, *P* < e-13 for all pairwise comparisons, Wilcoxon rank sum test, two-tailed). These results show that ISF has little effect on relative quantification of peptide precursors.

So far, our analyses have been at the peptide precursor level but most applications in proteomics rely on peptide or protein level analyses, in which data of peptide precursors that only differ in charge or even PTMs are combined to improve quantification and reduce redundancy. Since ISF only had a minor impact on relative peptide precursor quantification, we tested if assigning in-source fragments to their parental peptide groups or protein groups provide an advantage, e.g. by reducing errors, or if in-source fragments should simply be filtered out.

First, we used network analysis to group in-source fragments with their respective ISF parents, and calculated peptide quantities by summing all peptide precursors intensities of each peptide group. Subsequently, we compared the peptide groups of ISF parents with and without added in-source fragments as well as all the peptide groups unaffected by ISF with each other. The CVs of peptide groups that contain in-source fragments were smaller than the same peptide groups without in-source fragments, although the difference was minor (Fig. 6.d, *P* = 0.013, Wilcoxon rank sum test, two-tailed). The absolute relative errors showed that parent peptides (without merging with in-source fragments) much better followed the theoretical fold changes than non-ISF peptides (Fig. 6.d). However, these data also revealed that merging in-source fragment data with their parental peptide groups worsens quantification (*P* = 2.6e-14, Wilcoxon rank sum test, two-tailed) but again the effect is small.

Second, we calculated protein-level quantities by summing up all peptide precursors that belong to a certain protein, once with in-source fragments and once without them. Comparing the CVs and relative errors revealed that in-source fragments had close to no effect on the protein-level quantities (Fig. 6.e, *P* > 0.4, Wilcoxon rank sum test, two-tailed).

Taken together, these results show that ISF has little effect on relative quantification, that in-source fragments and parent peptides are among those peptides that are usable for relative quantification, and that ISF parents were the best peptides for quantification. These findings are likely because in-source peptides have typically higher intensities than non-ISF peptides. Regrouping in-source fragments with their parent peptides or including them in protein groups also had little effect, with a small tendency to reduce CVs of the new peptide groups but at the cost of distorting relative quantities across samples. In sum, while in-source fragments present no major challenge for quantitative analyses in general, they are only a proxy of their parent peptides, with slightly worse quantitative performance (CVs and relative errors), and we therefore recommend filtering them out.

### Impact of ISF on immunopeptidomics

Since ISF of tryptic peptides produces mostly peptides with semi-tryptic sequences (Supplementary Fig. 4), we wondered how ISF would impact peptide-centric proteomic approaches relying on relaxed trypsin specificity searches, such as immunopeptidomics. During the immune response, peptides produced by proteolytic *in vivo*-processes are presented to killer T cells as antigens^25,53^. Identifying those antigens is crucial for a better understanding of immune responses but also for the development of new medical treatments. Misidentification of an in-source fragment as true biological antigen can thus have costly consequences. The peptide antigens show narrow length distributions, typically between 9 and 14 amino acids, depending on the peptide class, and they do not show specific terminal amino acids^54^. Therefore, immunopeptidomics requires unspecific searches for peptide identification, which causes an exponential increase of the search space and increases the risk of confusing in-source fragments with biologically relevant peptides.

Here, we used four datasets from the literature (PXD022950, PXD051490, PXD054417, and PXD05880)^55–58^ to assess the impact of ISF on immunopeptidomics. Across the datasets, we observed large variation in the number of in-source fragments (Fig. 7.a), and similarly to our previous results, most ISFs occurred at the N-terminus (Fig. 7.b). As expected, the length distribution of peptides from immunopeptidomes was much narrower than in our data of trypsin-digested proteomes, but in-source fragments were again typically 1 to 2 amino acids shorter than parent peptides (Fig. 7.c). In the datasets PXD058880 and PXD022950, in-source fragments made up 16.6 and 36.8 % of all peptides below 9 amino acids (2.6 and 2.3 % by intensity) but only 0.07 and 0.92 % of longer peptides (0.04 and 0.8 % by intensity), respectively. The other datasets that we analyzed (PXD054417 and PXD05149) had much less ISF; the fragment identifications were again higher for peptides below 9 amino acids (0.53 and 0.90 % by identifications, 0.003 and 0.17 % by intensity) than for longer peptides (0.11 and 0.07 % by identifications, 0.06 and 0.01 % by intensity).

**Fig. 7.**
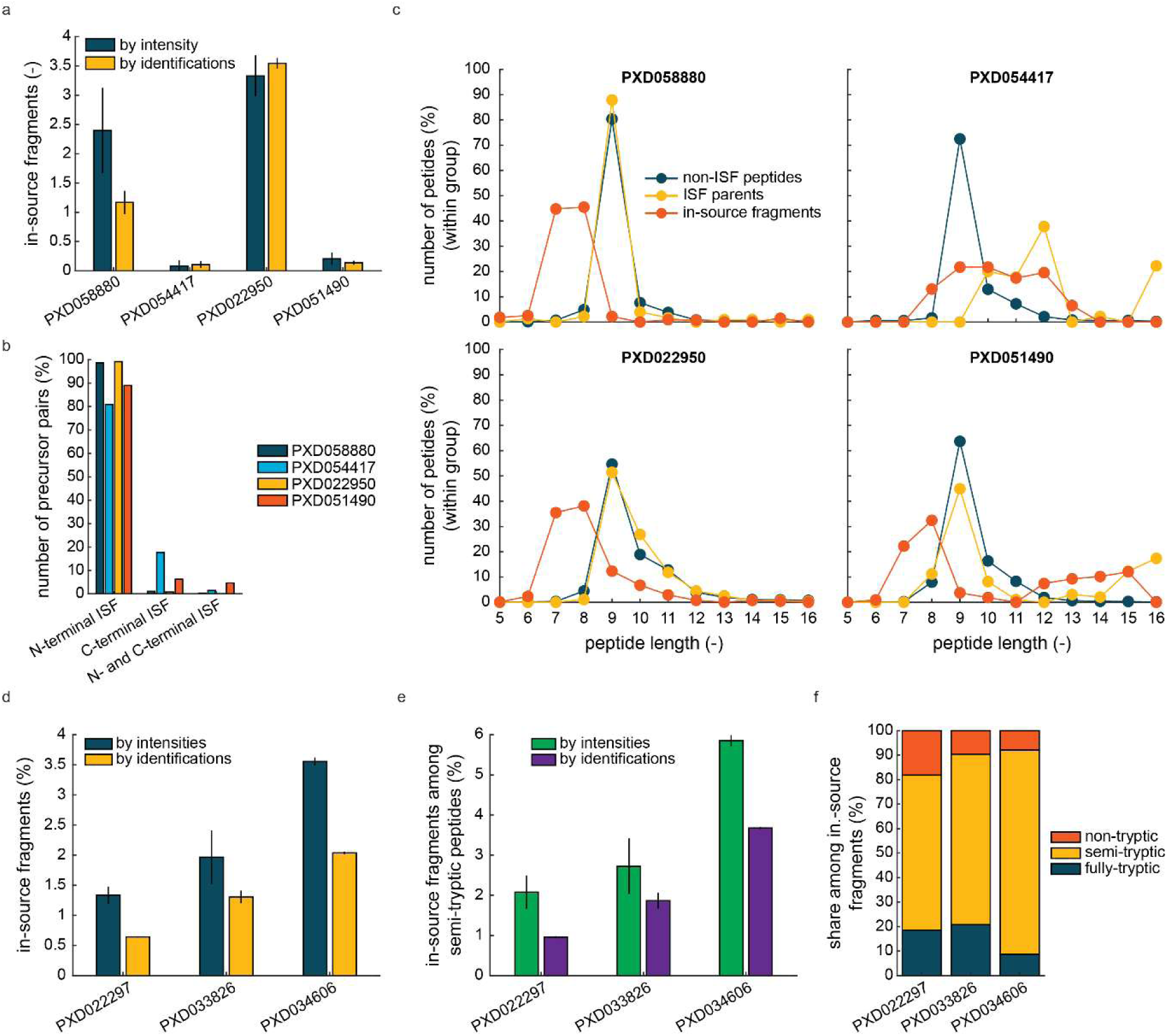
ISF in immunopeptidomics and limited proteolysis datasets. **a,** Bar plot showing the average percentage of in-source fragments in immunopeptidomics datasets (PXD058880: n = 12, PXD054417: n = 12, PXD022950: n = 6, PXD051490: n = 12), by peptide intensities and numbers of identifications. **b,** Bar plot showing the number of precursor pairs with evidence of ISF (per dataset in A) (%) that feature N-terminal ISF, C-terminal ISF, or ISF at both termini. **c,** Dot plots displaying the number of peptides (%, normalized per group) of different lengths (number of amino acids) in each dataset of A. Non-ISF peptides are indicated in blue (ISF parents: yellow, in-source fragments: red). **d,** Bar graph showing the mean fraction of in-source fragments within each indicated LiP dataset (yellow bars: % by intensity, blue bars: % by the number of identifications) (n = 3). Data results from trypsin unspecific peptide searches. **e,** Bar plot showing the percentage of in-source fragments among semi-tryptic peptides (green bars: % by intensity, violet bars: % by identifications) in the indicated LiP datasets. **f,** Bar plot showing the percentage of in-source fragments by their apparent trypticity in the indicated LiP datasets. Red bars indicate in-source fragments that appear as non-tryptic peptides (yellow bars: semi-tryptic in-source fragments, blue bars: fully-tryptic in-source fragments).

These results show that, in immunopeptidomics based on HLA-I peptides, ISF is likely to impact peptides shorter than 9 amino acids. Therefore, a simple strategy to mitigate this would be to exclude peptides shorter than 9 amino acids before further analyses. Alternatively or in addition, application of our RT-based detection algorithm to immunopeptidomics data will enable identifying in-source fragments of any length that contaminate the pool of biologically relevant peptides.

### In-source fragmentation in limited proteolysis data

Limited proteolysis coupled with mass spectrometry (LiP-MS) is another peptide-centric proteomic approach, which enables the global analysis of protein structural changes in proteome extracts^26^. It relies on proteases with broad specificity that cleave proteins for a brief period of time such that structural properties govern cleavage events. These primary cleavages can produce large protein fragments that may not be directly amenable for MS and are thus further digested by trypsin under denaturing conditions in a second step. LiP-MS therefore intrinsically features many more semi-tryptic peptides than other proteomic datasets. As most in-source fragments appear as semi-tryptic peptides based on our analyses, we asked how many of the identified semi-tryptic peptides in a typical LiP experiment were due to ISF and therefore could mistakenly be identified as structurally relevant peptides.

We used three published LiP datasets to study this (PXD022297, PXD033826, and PXD034606)^59–61^ and found that 0.6 % to 2.0 % of all identified peptides were due to ISF (1.3 to 3.6 % of the total intensity) (Fig. 7.d). However, the in-source fragments made up only 0.9 to 3.7 % of the semi-tryptic peptides (2.1 to 5.9 % by intensity), showing that most semi-tryptic peptides were indeed true-positives. In these LiP datasets, only 7.9 to 18.1 % of the in-source fragments appeared were non-tryptic (Fig. 7.f), corresponding to 0.08 – 0.12 % of all identifications (0.07 – 0.14 % by intensity).

These results show that ISF had only little impact on LiP datasets as the vast majority of semi-tryptic peptides were indeed caused by limited proteolysis and not by ISF. While the inherently high amounts of semi-tryptic peptides in LiP data did yield a small number of in-source fragments, detection of these peptides requires unspecific searches, which are typically not performed for standard LiP datasets. Overall, our data indicates that ISF is unlikely to be an issue for LiP-MS analyses.

### Current peptide-centric DIA analyses underestimate the extent of ISF

Our analyses so far focused on DIA datasets because DIA offers higher confidence than DDA in assessing co-elution of peptides and therefore in detecting ISF. DIA data however come with an important caveat: most peptide-centric DIA algorithms assume that a peptide elutes as a single peak at a specific retention time. However, if ISF occurs, it is possible that a peptide has more than one peak because it could stem from ISF in addition to a biological source that gives rise to a genuine peptide. Therefore, current search engines will either miss the identification of a genuine peptide or underrepresent ISF in cases in which a peptide has multiple peaks (Supplementary Fig. 11.a and b). In contrast to current DIA analysis approaches, DDA analysis does not make any assumption regarding the elution of peptides but reports all RTs at which a sequence is identified, although RT determination can be imprecise due to dynamic peak exclusion settings (Supplementary Fig. 11.C).

To understand the extent to which current “single-peak” peptide-centric DIA algorithms underestimate ISF, we analyzed DIA datasets using an approach that relies on modified DDA libraries^62^ (also see Methods part M.5). In brief, peptides that were identified in multiple DDA windows (each two or more minutes apart) were assigned to different bins of spectral assays using unique identifiers. Searches of DIA data with such modified spectral libraries would thus retain assays of the same peptide that are separated by RT and allow analysis of multiple peaks. We used this strategy to generate a modified spectral library from the DDA files acquired for the LiP dataset PXD022297^59^, which contains many genuine semi-tryptic peptides due to limited proteolysis. The library had 44,046 peptide precursor assays and included 2,961 (6.7 %) assays of precursors that were present in more than one RT window. Next, we used this library to search the DIA data from the same dataset and compared the library-based peptide peak identifications to those obtained by a “direct” peptide search of the DIA data, which did not rely on a spectral library from DDA. This revealed that the library-free DIA extraction identified 1,505 (57 %) precursors of the 2,961 assays with multiple peaks from our modified spectral library at the expected RT, misidentified 940 assays at a different RT, and completely missed 211 assays. Using the ISF parent and fragment annotations of the library-free DIA extraction showed that the (direct) peptide-centric DIA analysis underestimated the number of ISF peptides by 218 and the number of genuine peptide sequences by 933.

Using a spectral library for DIA extraction introduces the usual biases of DDA measurements, such as undersampling low abundant signals or underrepresenting singly charged precursors. Therefore, our results on the multiple peak extractions still present an underestimation of ISF and the true number of cases in which a peptide sequence occurs multiple times in a gradient. However, this analysis shows that current peptide-centric DIA analysis tools that only account for a single RT per peptide sequence systematically underestimate both the extent of ISF as well as the number of genuine peptides, which are not created by ISF. These results thus encourage the development of peptide-centric algorithms that account for multiple peaks.

## Discussion

It is known that ISF affects peptides and can be a source of peptides with non-tryptic termini in bottom-up proteomics data^21^. The proteomics field has evolved in past years, with an increasing use of peptide-centric proteomics approaches like structural proteomics and immunopeptidomics that rely on semi- and non-tryptic peptides. Further, DIA methods that enable detection of low intensity *m/z* features, as opposed to DDA methods that likely miss many low intensity fragments, have found mainstream adaptation, and the development of the new mass spectrometers with vastly increased sensitivity has introduced new instrument geometries and designs that could affect the prevalence of ISF. It is therefore timely to conduct an assessment of ISF in the context of DIA proteomics, with a particular focus on peptide-based approaches.

We described an approach to re-annotate peptides based on RT patterns. The novelty of our approach is to use intra-sample information from peptides with multiple charge states to estimate natural distributions of ΔRT values and, based on these, determine ΔRT cutoffs for ISF detection. This approach can therefore account for differences in LC performance between runs, batches, or HPLCs. In the future, our ISF detection could even be further improved by using dynamic ΔRT cutoffs that depend on the RT of the peptide peaks. This could address dynamically varying ΔRT value distributions in non-linear gradients.

We observed that ISF can affect a large share of a proteomics dataset acquired by DIA and account for more than 50 % of the detected semi-tryptic peptides. We also found that the numbers of detected in-source fragments anticorrelate with sample complexity; this has implications for MS approaches such as crosslinking MS, hydrogen/deuterium exchange mass spectrometry (HDX-MS), affinity purification coupled to MS, or (co-)fractionation MS that often rely on the analysis of low complexity samples^63–66^. These observations match well with previous reports based on DDA data^21^. Further, our analysis of the intensity distributions showed that the observed ISF parents often had very high intensities resulting also in fragments with higher intensities than of non-ISF peptides. This observation could be the result of a bias of the search engine against in-source fragments with low intensities.

Generally, we observed a large variation in the amount of detected ISF among different datasets, which can be explained by different samples complexities, instruments, and chosen parameters, all of which we have shown here to have an impact on the prevalence of ISF. The combinatorial effect of all these factors makes it difficult to estimate whether ISF could be a problem for a particular sample or dataset and, considering that ISF can in some cases account for more than 30 % of the peptide identifications, we suggest that ISF products should be detected and annotated as a routine data analysis step.

The current peptide-centric DIA analysis tools assume that a peptide sequence occurs only once across a chromatogram. Using a RT-based splitting strategy to generate spectral libraries from DDA data, we were able to analyze peptides occurring multiple times across the LC gradient in DIA data. Our analysis showed that both the extent of ISF and the number genuine, non-ISF peptides were underestimated depending on the peak selection. Underestimating ISF might not present a problem for typical applications but, whenever an in-source fragment is identified, it can come at the cost of losing an identification of a genuine peptide that shares its sequence with the in-source fragment. With current peptide search strategies for DIA data, ISF can thus mask the identification of genuine peptides and lead to loss of information. Furthermore, for peptides with high signals, there is a reasonable chance that an in-source fragment can be observed in the data. This effect could be used in future peptide search engines to improve peptide identification and estimation of false-discovery rates while increasing overall annotations of proteomics data. In immunopeptidomics, peptides are not generated by *in vitro* digestion and are often identified by unspecific peptide searches. Therefore, immunopeptidomics data could be much more prone to interference by ISF, which often generates peptides with non-tryptic termini. Our results show that in immunopeptidomics data with HLA-I peptides, most in-source fragments are shorter than 9 amino acids and can account for more than a third of all peptides of such short length. This raises the question of how many of the immunopeptides shorter than 9 amino acids are truly authentic peptides because current approaches still underestimate how many of the detected peptides are in-source fragments. We recommend using our ISF detection to minimize the risk of falsely identifying peptides. Alternatively, or in addition, immunopeptides smaller than 9 amino acids can be filtered out to strongly reduce the impact of ISF on the data.

As in-source fragments have the same RT as their parent peptides, the RT of the in-source fragment does not correctly reflect the peptide separation by chromatography and is different from the RT of a non-ISF peptide with the same amino acid sequence. Peptide ISF can therefore have an impact on the RT realignment during peptide searches. Depending on the MS approach, the strength of this effect is likely to vary, and will be more impactful in cases where a few reference peptides are chosen for realignment, like analyses by single reaction monitoring (SRM), or when the prevalence of ISF is particularly high.

We observed little PTM ISF for phosphorylation, oxidation of methionine, and N-terminal acetylation, but we cannot exclude that other PTMs such as glycosylation, which is known to be prone to ISF^67^, might be affected to a much larger extent. Some PTMs might even be so labile that we only detect the in-source fragments, which would require exceptionally mild ionization and ion routing approaches to study them. Sequence-altering ISF of parent peptides that carry a PTM could also explain some of the detected semi-tryptic peptides in typical trypsin-digested bottom-up proteomics samples, which have not been associated with an ISF parent so far, because many peptides with PTMs are very challenging to detect in non-enriched datasets by current search engines. Other cases of ISF that have not been addressed so far are those in which fragmentation occurs not at peptide bonds (between C1 and N2) or at PTM-peptide bonds but rather at side chains. Detecting these cases presents a major challenge but potentially could explain many of the detected but uncharacterized *m/z* features in proteomics data and help separate these from more biologically meaningful uncharacterized portions of the proteome^68^. Moreover, in-source chemical reactions that add to the mass of a molecule have been observed for metabolites before at a high prevalence^13^ and, in principal, could also affect peptides. Taken together, this poses the question: how many of the *m/z* features in a proteomics dataset that are unannotated can be explained by ISF or in-source reactions? Answering this question will result in new peptide groups without annotation, for which the mode of fragmentation or reaction could provide insights into the identity of the parent peptides.

In summary, in-source fragments inflate the number of identifications and, especially in low complexity samples, can make up most of the signal and lead to data misinterpretation. Further, since ISF varies substantially dependent on numerous parameters, predicting the impact of these fragments *a priori* in a given analysis is challenging. Based on our findings in this study, we thus conclude that ISF should be identified by default in current proteomics approaches and the in-source fragments filtered out bioinformatically.

## Methods

### ***M.1*** Sample preparation for MS-based proteomics

#### M.1.1 Saccharomyces cerevisiae S288c whole-proteome sample

*S. cerevisiae* S288c (ATCC #204508) was streaked from cryo stock onto an YPD agar plate and incubated at room temperature. After single colonies became visible, the plate was stored at 4 °C. 25 mL synthetic defined (SD) medium containing 20 g/L D-glucose (Merck, Sigma-Aldrich #G7021), 5 g/L ammonium sulfate (Fluka #09982), and yeast base (Merck, Millipore #Y1251) was inoculated from a single colony and incubate in 100 mL Erlenmeyer flasks at 30 °C under shaking of 200 rpm for circa 20 h. 40 mL SD medium (2 w/v-% glucose) in 500 mL Erlenmeyer flasks was inoculated from the previous 25 mL-culture to a start optical density (OD at 600 nm) of 0.004 and incubated at 30 °C under shaking at 200 rpm. 300 mL SD medium (2 w/v-% glucose) in 2 L Erlenmeyer flasks were inoculated from the previous 40 mL-culture to a start OD of 0.08 and incubated at 30 °C under shaking at 200 rpm. The OD was measured regularly to calculate growth rates and exclude growth defects. At a final OD of circa 0.6, samples for proteomics were collected using a method that is analogous to a phosphoproteomics sampling approach^45^ and relies on quenching enzyme activity by trichloroacetic acid (TCA): 275 mL of culture were transferred to 500 mL harvesting tubes on ice. 18.3 mL 4 °C-cold 100 w/v-% TCA (final concentration 6.2 % TCA; Merck, Sigma-Aldrich, #91228) was added. The cell suspension was incubated for 10 min on ice and, subsequently, centrifuged for 5 min at 4 °C and 3428 g. The supernatant was discarded, and the cell pellet resuspended in 40 mL 4 °C-cold acetone (Merck, Supelco, #1.00014). The suspension was transferred to 50 mL reaction tubes and centrifuged for 5 min at 4 °C and 3,428 g. The supernatant was discarded, the cell pellet resuspended in 40 mL 4 °C-cold acetone, and the suspension centrifuged for 5 min at 4 °C and 3,428 g. The supernatant was discarded, and the cell pellet resuspended in 5 mL 4 °C-cold acetone. The cell suspensions of 4 independent biological replicates (= 4 different colonies) were pooled and distributed to 1.5 mL screw cap-reaction tubes. After 5 min of centrifugation at 3,400 g and 4 °C, the supernatant was discarded, and the remaining cell pellets flash frozen in liquid nitrogen and stored at -80 °C. Cell pellets were resuspended in 500 µL lysis buffer, which was 8 M urea (Merck, Empure Essential, #1.08486.1000) 100 mM ammonium bicarbonate (Merck, Sigma-Aldrich, #A6141) at pH 7.8. After adding an equal volume of glass beads (0.5 mm diameter, Merck, Sigma-Aldrich # G8772), cells were lysed at 4 °C using 8 cycles of bead beating at 6.5 m/s for 30 s with 200 s pause. After bead removal, lysates were centrifuged for 15 min at 17,000 g and 4 °C, and all supernatants pooled for further processing. After tryptic digest and peptide clean-up, the reference yeast sample was obtained by mixing 800 µL of the *S. cerevisiae* S288c sample with 200 µL of the *mix of 8 proteins sample*.

### M.1.2 Escherichia coli BW25113 whole-proteome sample

*E. coli* BW25113 (DSMZ, #27469) was streaked from cryo stock onto a LB agar plate and incubated at 37 °C overnight. The plate was stored at 4 °C. 25 mL M9 minimal medium was inoculated from a single colony and incubate in 100 mL Erlenmeyer flasks at 37 °C under shaking of 200 rpm overnight. M9 minimal medium contained: 5 g/L D-glucose (Merck, Sigma-Aldrich, #G7021), 6 g/L Na_2_HPO_4_ (Carl Roth, #P030.2), 3 g/L KH_2_PO_4_ (Carl Roth, #3904.1), 0.5 g/L NaCl (Carl Roth, #9265.1), 1.5 g/L (NH_4_)_2_SO_4_ (Merck, Sigma-Aldrich, #A3920), 1.8 mg/L ZnSO_4_x 7 H_2_O (Fluka, #96500), 1.2 mg/L CuCl_2_ x 2 H_2_O (Fluka, #61174), 1.2 mg/mL MnSO_4_ x H_2_O (Merck, Sigma-Aldrich, #M7631), 1.8 mg/L CoCl_2_ x 6 H_2_O (Riedel de Haën, #12914), 2.8 mM thiamine-HCl (Merck, Sigma-Aldrich, #T4625), 1mM MgSO_4_ x 7 H_2_O (Merck, Sigma-Aldrich, #63138), 100 µM CaCl_2_ x 2 H_2_O (Merck, Sigma-Aldrich, #C8106), and 100 µM FeCl_3_ x 6 H_2_O (Merck, Sigma-Aldrich, #31232). 300 mL M9 minimal medium (0.5 w/v-% glucose) in 2 L Erlenmeyer flasks were inoculated from the previous 25 mL-culture to a start OD of circa 0.004 and incubated at 37 °C under shaking at 200 rpm. The OD was measured regularly to calculate growth rates and exclude growth defects. At a final OD of circa 0.6, samples for proteomics were collected: 275 mL of culture were transferred to 500 mL harvesting tubes on ice. 18.3 mL 4 °C-cold 100 w/v-% TCA was added. The cell suspension was incubated for 10 min on ice and, subsequently, centrifuged for 5 min at 4 °C and 3,428 g. The supernatant was discarded, and the cell pellet resuspended in 40 mL 4 °C-cold acetone. The suspension was transferred to 50 mL reaction tubes and centrifuged for 5 min at 4 °C and 3,428 g. The supernatant was discarded, the cell pellet resuspended in 40 mL 4 °C-cold acetone, and the suspension centrifuged for 5 min at 4 °C and 3,428 g. The supernatant was discarded, and the cell pellet resuspended in 5 mL 4 °C-cold acetone (Merck, Supelco, #1.00014).

The cell suspensions of 4 independent biological replicates (= 4 different colonies) were pooled and distributed to 1.5 mL screw cap-reaction tubes. After 5 min of centrifugation at 3,400 g and 4 °C, the supernatant was discarded and remaining cell pellets flash frozen in liquid nitrogen. Samples were stored at -80 °C. Cells were lysed like *S. cerevisiae* S288c samples.

### M.1.3 Homo sapiens HEK-293 whole-proteome sample

*H. sapiens* HEK-293 cells (ATCC, #CRL-1573) were thawed from cryo stock and transferred to a cell culture-grade, sterile, 10 cm-diameter petri dish. 10 mL Dulbecco’s Modified Eagle Medium (DMEM, Thermo Fisher Scientific, Gibco #41965039) containing 10 v/v-% heat inactivated foetal calf serum (FCS, BioConcept, #2-01F16-I) and 1 v/v-% penicillin-streptomycin solution (P/S, 10,000 U/mL, Thermo Fisher Scientific, Gibco #15140122) were added, and cells incubated for 3 days at 37 °C, 5 %CO_2_, and a relative humidity of 85 %. The supernatant was removed, 10 mL phosphate buffered saline solution (PBS, pH 7.4, Thermo Fisher Scientific, Gibco #10010015) added to wash surface adherent cells. The supernatant was removed and 1 mL 0.25 % Trypsin-EDTA solution (Thermo Fisher Scientific, Gibco #25200056) added. 24 mL of fresh medium were added to a 15 cm-diameter petri dish, trypsin-suspended cells transferred to the plate with fresh medium, and cells incubated for 4 days. The cell culture was tested negative for mycoplasma contamination by the MycoGenie Rapid Mycoplasma Detection Kit (AssayGenie, #MORV0011-50). The supernatant was removed and cells washed with 25 mL PBS. 3 mL 0.25 % Trypsin-EDTA solution added, and 5 new plates each with 23 mL fresh medium prepared. 8 mL medium were added to the trypsinated culture, each 2 mL of the cell suspension added to a new plate, and cells incubated for 2 days. Proteomics samples were taken by transferring the plates onto ice, remove the supernatant, and, for each plate, adding 10 mL 4 °C-cold PBS containing 1 mM EDTA (Merck, Sigma-Aldrich, #03677). The cell suspension was transferred to 15 mL reaction tubes and centrifuged for 4 min at 300 g and 4 °C. The supernatant was removed, and cells resuspended in 1 mL of the 4 °C-cold PBS-EDTA solution. The cell suspensions from 4 plates were pooled, distributed to 2 mL reaction tubes, and centrifuged for 4 min at 300 g and 4 °C. The supernatant was discarded, and cell pellets flash frozen in liquid nitrogen and stored at -80 °C. Cell pellets were resuspended in 500 µL lysis buffer and lysed by vortexing twice for 30 s with a pause of at least 30 s on ice. Lysates were centrifuged for 15 min at 17.000 g and 4 °C. Supernatants were used for further processing.

### M.1.4 Mix of 8 proteins

Separate solutions of bovine catalase (Uniprot ID: P00432, Merck, Sigma-Aldrich, #C40), rabbit creatine kinase (P00563, Merck, Sigma-Aldrich, #C3755), rabbit fructose-bisphosphate aldolase A (P00883, Merck, Sigma-Aldrich, #A2714), bovine lactoferrin (P24627, Merck, Sigma-Aldrich, #L9507), chicken ovotransferrin (P02789, Fluka, #27695), rabbit pyruvat kinase (P11974, Merck, Sigma-Aldrich, #P9136), bovine serotransferrin (Q29443, Merck, Sigma-Aldrich, #T1408), and bovine serum albumin (P02769, Merck, Sigma-Aldrich, #A7638) were prepared at 1 µg protein/µL in 8 M urea 100 mM ammonium bicarbonate. After tryptic digestion and sample clean-up, peptides from the individual proteins were pooled together yielding the *mix of 8 proteins sample*.

### M.1.4 Tryptic digest and sample clean-up

Protein concentrations in cell lysates were measured using a bicinchoninic acid-based assay (Thermo Scientific, Pierce BCA Protein Assay Kit, #23225). At a protein concentration of 1 µg/µL, disulfide bonds were first reduced by incubation with 5 mM tris(2-carboxyethyl)phosphin - hydrochlorid (TCEP, Merck, Sigma-Aldrich, #C4706) for 30 min at 37 °C under 200 rpm of shaking and second alkylated by incubation with 12 mM iodoacetamide (IAA, Merck, Sigma-Aldrich, #I1149) for 15 min in the dark at 37 °C under 200 rpm of shaking. Samples were diluted with 100 mM ammonium bicarbonate (Merck, Sigma-Aldrich, #A6141) to a final urea concentration of 1 M. Sequencing-grade trypsin (Promega, #V5111) was added at a protease to protein ratio of 1:100, and samples incubated overnight at 37 °C. The digestion was stopped with formic acid (FA, Merck, Sigma-Aldrich, #33015) at a final concentration of 2 %. Samples were desalted using C18 columns (Waters Sep-Pak Vac 3cc, 500 ng). After washing the columns with 2.5 mL methanol, twice with 2.5 mL Buffer B (50 % 0.1 % FA), and thrice with 2.5 mL Buffer A (0.1 % FA), samples were loaded onto the columns, and columns washed thrice with 2.5 mL Buffer A. Peptides were eluted with 2 mL Buffer B, dried by vacuum centrifugation at 40 °C, and resuspended in Buffer A at a concentration of ca. 1 µg/µL.

### M.1.5 Affinity purification samples

A *H. sapiens* HEK-293 cell line expressing the Strep-HA-tagged human kinase SRPK3 (Uniprot ID: Q9UPE1) under an inducible promoter was obtained from Varjosalo *et al.*,^69^ and cells seeded at a density of 10^7^ in 25 mL DMEM supplemented with 10 % FCS and 1 % P/S in a 15 cm dish and incubated overnight at 37 °C. Protein expression was induced with 1.3 μg/mL doxycycline for 24 hours. Per sample, cells from two 15 cm dishes were harvested by scraping off in 4 °C-cold PBS. Cells were washed once with PBS, and cell pellets flash frozen in liquid nitrogen. Pellets were lysed by resolubilizing in 2 mL HNN buffer (50 mM HEPES pH 8.0, 150 mM NaCl, 50 mM NaF) supplemented with 0.5 % IGEPAL, 400 nM Na_3_VO_4_, 1 mM phenylmethylsulfonyl fluoride, 0.2 % protease inhibitor cocktail (Merck, Sigma-Adrich, #P8340) and 0.5 μL/mL benzonase (Merck, Sigma-Aldrich, #1.01695). The lysate was incubated on ice for 10 min and centrifuged at 18,000 g for 20 min at 4 °C. The supernatant was added to 80 μL StrepTatcin Sepharose resin (IBA Lifesciences, #2-1201) and incubated on a rotary shaker for 60 min at 4 °C. The beads were loaded onto a 1 μm glass filter column and washed 3 times with 1 mL HNN buffer supplemented with 0.5 % IGEPAL and 400 nM Na_3_VO_4_, 3 times with 1 mL HNN buffer, and 3 times with 100 mM ammonium bicarbonate. Samples were eluted twice by incubation with 30 μL of 0.5 mM biotin in 100 mM ammonium bicarbonate for 15 min. To the samples, 60 μL 8 M urea in 100 mM ammonium bicarbonate for a final concentration of 4 M urea. Samples were reduced with 5 mM TCEP for 40 min at 37 °C and 200 rpm and alkylated with 40 mM IAA for 30 min at 30 °C in the dark, at 200 rpm. Samples were diluted with 100 mM ammonium bicarbonate to a urea concentration of 1 M. To each sample, 1 μg Lys-C and 1 μg trypsin were added, and samples were incubated overnight at 37 °C and 200 rpm. The digestion was stopped with 2 % formic acid. Samples were desalted using HNFR S18V desalting plates (Nest Group). The resin was activated using 200 μL methanol and washed 3 times with 200 μL Buffer B. The resin was equilibrated 3 times with 200 μL Buffer A. Samples were loaded and washed 3 times with 200 μL Buffer A. Samples were eluted twice with 100 μL Buffer B and dried at 40 °C in a vacuum centrifuge.

### M.1.6 Mix of 11 peptides sample

We used a commercially available mix of 11 peptides (Biognosys iRT Kit, Bruker Daltonics, #1816351).

### M.1.7 Phosphoproteomics samples

Phosphopeptides of *S. cerevisiae* S288c samples were enriched based on a protocol described in Bodenmiller *et al.* ^45^. Between 0.5 and 1 mg of desalted peptides were incubated in a rotary shaker for 1 h with 1.25 mg of TiO_2_ resin (GL Sciences, Japan), preequilibrated twice with 500 µL of methanol, and twice with 500 µL of a solution saturated with phthalic acid (Merck, Sigma-Aldrich, #402915). Peptides bound to the TiO_2_ resin were washed twice with 500 µL phthalic acid solution, twice with 80 % acetonitrile 0.1% formic acid solution, and finally twice with 0.1% formic acid. The phosphopeptides were eluted from the beads twice with 150 µl of 0.3 M ammonium hydroxide at pH 10.5 and immediately acidified with 50 µL 5% trifluoroacetic acid to reach about pH 2.0. The enriched phosphopeptides were desalted on microspin columns (The Nest Group, USA), dried using a vacuum centrifuge, and resolubilized in 10 µl of 0.1 % formic acid.

### ***M.2*** LC-MS

If not stated otherwise, an EASY-nLC 1200 (Thermo Scientific) coupled to an Exploris 480 (Thermo Scientific) was used to measure proteomics samples in DIA mode. A sample volume corresponding to 1 µg of peptides was injected. A 40 cm x 0.75 µm (inner diameter) column (New Objective, 10 µm tip, PicoFrit, #PF360-75-10-N-5) packed with 3 μm C18 beads (Dr. Maisch, 120 Å pore size, 300 m² surface area, #Reprosil-Pur 120) was used for separating peptides over a linear 120 min gradient. Buffer A was 0.1 % FA, and Buffer B 50 % ACN 0.1 % FA. The gradient started at 3 % Buffer B and ended at 30 % Buffer B. The flow rate was 300 nL/min. Peptides were measured in positive ionization mode. The electrospray voltage was 2500 V, the ion transfer capillary temperature was 275 °C, and the funnel radiofrequency 50 % if not indicated else. To reduce contamination of the MS, the electrospray voltage was set to 1500 V during the initial 8 min of the method and to 0 V during column washes. The MS1 full scan range was from 350 to 1150 *m/z* at a resolution of 120,000. with an automatic gain control (AGC) target of 200 % and a maximum injection time of 264 ms. Precursors were fragmented in the higher-energy collisional dissociation (HCD) cell at 30 % relative collision energy. For DIA, 41 variable-size MS2 scan windows were used between 350 and 1,150 m/z, each at a resolution of 30,000, an AGC target of 200 %, a maximum injection time of 66 ms, and a window overlap of 1 Da.

For measurements with an Q Exactive Plus (Thermo Scientific), which was also coupled to an EASY-nLC 1000 (Thermo Scientific), the same parameters were used as for the Exploris 480, if possible. MS1 full scans differed by the resolution (70,000), the AGC target (3e6), and the maximum injection time (120 ms). DIA MS2 measurements differed by the resolution (35,000), AGC target (3e6), and the maximum injection time (60 ms).

For measurements with an Astral (Thermo Scientific) coupled to a Vanquish Neo LC system (Thermo Scientific), the flow rate was 400 nL/min, and a 41.8 min gradient was used. Starting with Buffer B at 3 %, buffer B was changed in linear steps to 32 % at 25.5 min, 45 % at 30.5 min, 95 % at 32.5 min, where it was kept constant for 8 min, and 3 % at 41.8 min. MS1 full scans were acquired between 350 and 1400 m/z with a resolution of 240,000. The AGC target was 5e6. For DIA, 524 fixed-size (2 Da) MS2 scan windows between 350 and 1400 m/z were used with a maximum injection time of 3 ms, a relative HCD collision energy of 27 %, the RF lens at 40 %, and an AGC target of 5e4.

### ***M.3*** Peptide searches and data processing

Spectronaut 20 (Biognosys, Schlieren, Switzerland) was used to search DIA data for peptides using “directDIA”. The data was searched against different proteome databases appropriate to each sample and related FASTA-files were obtained from UniProt^70^. The enzyme specificity was set to semi-specific with cleavage rules for trypsin (or if applicable also for LysC). Peptide lengths between 7 and 52 amino acids, two missed cleavages, and a maximum of 5 variable modifications were permitted. The immunopeptidomics and LiP datasets were searched for peptides with unspecific cleavage sites and 5 to 16 amino acids in length. N-Acetylation at protein N-termini and oxidation of methionine were set as variable modifications, and carbamidomethylation of cysteine as a fixed modification. Searches for phosphopeptides included phosphorylation of serine, threonine, and tyrosine. Data from the parameter test (also see Fig. 4) was searched twice: (1) all samples were searched individually to obtain an accurate representation of the number of peptide precursor identifications, and (2) all samples were searched together to obtain accurate quantitative data (intensities). Supplementary Table 2 provides an overview of which samples of the literature datasets (also see Supplementary Tab. 1) were used in this study and included in peptide searches.

Data from peptide searches were analyzed with MATLAB (R2023b, 23.2.0.2409890). Venn diagrams were created using the MATLAB function “venn”^71^. The data was analyzed at the peptide precursor level if not specified else. Precursors with identification Q-values greater than 10^-3^ and precursors without quantities or “FGMS2RawQuantity” smaller than 2 were filtered out, except for the algorithm comparison (also see Supplementary Fig. 3) for which no filter was applied.

### ***M.4*** ISF detection

ISF were detected in each sample separately. First, all peptide precursors were paired up against each other, for which one precursor’s amino acid sequence contained the sequence from the other precursor. Then, looping over all precursors and their pairings from small to large (length), ΔRTs were calculated by subtracting the peak apex retention times of the smaller peptide from the larger peptide. Subsequently, ΔRT cutoffs were determined by fitting the model function (EQ1) to the ΔRT distribution of precursor pairs that only differ in charge. The function consisted of two components. One of which was a normal distribution with mean value *µ* and standard deviation *σ* that was scaled with parameter *a*. The other component was the uniform distribution *k_0_*. ΔRT cutoff was the ΔRT, for which the absolute difference between the normal distribution and *k_0_* was smallest.

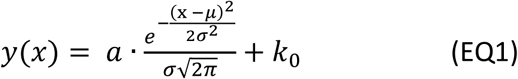

The distribution of measured ΔRTs was estimated ten times with different bin sizes and, accordingly, the model function fitted to each estimation using least square optimization. The final ΔRT cutoff was the median across the ten ΔRT cutoffs from the ten individual fits. If the number of datapoints of false-positive precursor pairs that only differ in charge was insufficient to model the uniform distribution, the ΔRT cutoffs could not be accurately determined and typically became very large. In such cases, in which our model-based approach yields a ΔRT cutoff larger than 0.15 min, the full width at half maximum (FWHM) of the normal distribution was used. In extreme cases, in which the data was even insufficient to model the normal distribution of true positives, a ΔRT cutoff of 0.1 min was taken. The lower boundary of the ΔRT cutoff was the smallest ΔRT value of the respective sample.

### ***M.5*** Multiple peak extraction from DIA data

To analyze peptide precursors with multiple peaks in DIA data, we followed a previously published strategy^62^. If the same peptide sequence is identified in DDA beyond a certain time span, which is typically longer than the average elution peak width (e.g. 2 min, which we used here), the peptide-spectrum matches were binned into separate spectral libraries. Subsequently, for each of the peptides, a local average retention time and consensus MS/MS fragmentation spectrum were calculated. To force the DIA analysis software to extract multiple identical peptide sequences at different retention times, we “anonymized” the peptides with a set of characters that cannot be properly parsed by the software (e.g. adding numbers or special characters). When faced with such non-parsable sequences, the search tool defers back to the only other information available to extract the DIA data: (1) the m/z and charge state of the precursor, (2) the *m/z*, charge states and relative intensities of the fragments, and (3) the peptide retention time. This strategy enabled us to estimate how often a peptide sequence could be identified multiple times in DIA file.

## Data and code availability

Proteomics data will be made available on PRIDE and relevant MATLAB code will be provided on GitHub and Zenodo upon publication. The source data of figures will be provided as supplementary information upon publication.

## Acknowledgments

We thank Oliver Bernhardt, Tejas Gandhi, Roland Bruderer, and Monika Pepelnjak for discussions. We thank Biognosys for early access to the ISF detection feature of Spectronaut 20. P. P. was funded by the Promedica Stiftung (2025-0022/M) and by an ETH Zurich Grant (25-2 ETH-027).

## Author contributions

*Thorben Schramm:* Conceptualization, Methodology, Software, Validation, Formal analysis, Investigation, Resources, Data Curation, Writing - Original Draft, Writing - Review & Editing, Visualization, Project administration

*Ludovic Gillet:* Conceptualization, Methodology, Software, Validation, Formal analysis, Investigation, Resources, Writing - Original Draft, Writing - Review & Editing

*Viviane Reber:* Resources

*Natalie de Souza*: Writing - Review & Editing

*Matthias Gstaiger:* Supervision

*Paola Picotti:* Conceptualization, Writing - Original Draft, Writing - Review & Editing, Supervision, Project administration, Funding acquisition

## References

1. Alseekh, S. et al. Mass spectrometry-based metabolomics: a guide for annotation, quantification and best reporting practices. Nat. Methods 18, 747–756 (2021).

2. Collins, S. L., Koo, I., Peters, J. M., Smith, P. B. & Patterson, A. D. Current Challenges and Recent Developments in Mass Spectrometry–Based Metabolomics. Annu. Rev. Anal. Chem. 14, 467–487 (2021).

3. Guo, T., Steen, J. A. & Mann, M. Mass-spectrometry-based proteomics: from single cells to clinical applications. Nature 638, 901–911 (2025).

4. Aebersold, R. & Mann, M. Mass-spectrometric exploration of proteome structure and function. Nature 537, 347–355 (2016).

5. Thomas, S. N., French, D., Jannetto, P. J., Rappold, B. A. & Clarke, W. A. Liquid chromatography–tandem mass spectrometry for clinical diagnostics. Nat. Rev. Methods Primer 2, 96 (2022).

6. Glish, G. L. & Vachet, R. W. The basics of mass spectrometry in the twenty-first century. Nat. Rev. Drug Discov. 2, 140–150 (2003).

7. Yamashita, M. & Fenn, J. B. Electrospray ion source. Another variation on the free-jet theme. J. Phys. Chem. 88, 4451–4459 (1984).

8. Prabhu, G. R. D., Williams, E. R., Wilm, M. & Urban, P. L. Mass spectrometry using electrospray ionization. Nat. Rev. Methods Primer 3, 23 (2023).

9. Daniel, J. M., Friess, S. D., Rajagopalan, S., Wendt, S. & Zenobi, R. Quantitative determination of noncovalent binding interactions using soft ionization mass spectrometry. Int. J. Mass Spectrom. 216, 1–27 (2002).

10. Guo, J., Shen, S., Xing, S., Yu, H. & Huan, T. ISFrag: De Novo Recognition of In-Source Fragments for Liquid Chromatography–Mass Spectrometry Data. Anal. Chem. 93, 10243–10250 (2021).

11. Xu, Y.-F., Lu, W. & Rabinowitz, J. D. Avoiding Misannotation of In-Source Fragmentation Products as Cellular Metabolites in Liquid Chromatography–Mass Spectrometry-Based Metabolomics. Anal. Chem. 87, 2273–2281 (2015).

12. Domingo-Almenara, X. et al. Autonomous METLIN-Guided In-source Fragment Annotation for Untargeted Metabolomics. Anal. Chem. 91, 3246–3253 (2019).

13. Farke, N., Schramm, T., Verhülsdonk, A., Rapp, J. & Link, H. Systematic analysis of in-source modifications of primary metabolites during flow-injection time-of-flight mass spectrometry. Anal. Biochem. 664, 115036 (2023).

14. Chen, L. et al. Widespread occurrence of in-source fragmentation in the analysis of natural compounds by liquid chromatography–electrospray ionization mass spectrometry. Rapid Commun. Mass Spectrom. 37, e9519 (2023).

15. Schmid, R. et al. Ion identity molecular networking for mass spectrometry-based metabolomics in the GNPS environment. Nat. Commun. 12, 3832 (2021).

16. Lu, W. et al. Improved Annotation of Untargeted Metabolomics Data through Buffer Modifications That Shift Adduct Mass and Intensity. Anal. Chem. 92, 11573–11581 (2020).

17. Giera, M., Aisporna, A., Uritboonthai, W. & Siuzdak, G. The hidden impact of in-source fragmentation in metabolic and chemical mass spectrometry data interpretation. Nat. Metab. 6, 1647–1648 (2024).

18. El Abiead, Y., et al. Discovery of metabolites prevails amid in-source fragmentation. Nat. Metab. 7, 435–437 (2025).

19. Senan, O. et al. CliqueMS: a computational tool for annotating in-source metabolite ions from LC-MS untargeted metabolomics data based on a coelution similarity network. Bioinformatics 35, 4089–4097 (2019).

20. Fuhrer, T., Heer, D., Begemann, B. & Zamboni, N. High-Throughput, Accurate Mass Metabolome Profiling of Cellular Extracts by Flow Injection–Time-of-Flight Mass Spectrometry. Anal. Chem. 83, 7074–7080 (2011).

21. Kim, J.-S., Monroe, M. E., Camp, D. G. I., Smith, R. D. & Qian, W.-J. In-Source Fragmentation and the Sources of Partially Tryptic Peptides in Shotgun Proteomics. J. Proteome Res. 12, 910–916 (2013).

22. Gillet, L. C. et al. Targeted Data Extraction of the MS/MS Spectra Generated by Data-independent Acquisition: A New Concept for Consistent and Accurate Proteome Analysis*. Mol. Cell. Proteomics 11, O111.016717 (2012).

23. Gillet, L. C., Leitner, A. & Aebersold, R. Mass Spectrometry Applied to Bottom-Up Proteomics: Entering the High-Throughput Era for Hypothesis Testing. Annu. Rev. Anal. Chem. 9, 449–472 (2016).

24. Ting, Y. S. et al. Peptide-Centric Proteome Analysis: An Alternative Strategy for the Analysis of Tandem Mass Spectrometry Data*. Mol. Cell. Proteomics 14, 2301–2307 (2015).

25. Chong, C., Coukos, G. & Bassani-Sternberg, M. Identification of tumor antigens with immunopeptidomics. Nat. Biotechnol. 40, 175–188 (2022).

26. Feng, Y. et al. Global analysis of protein structural changes in complex proteomes. Nat. Biotechnol. 32, 1036–1044 (2014).

27. Kalxdorf, M., Müller, T., Stegle, O. & Krijgsveld, J. IceR improves proteome coverage and data completeness in global and single-cell proteomics. Nat. Commun. 12, 4787 (2021).

28. Midha, M. K. et al. A comprehensive spectral assay library to quantify the Escherichia coli proteome by DIA/SWATH-MS. Sci. Data 7, 389 (2020).

29. Jayavelu, A. K. et al. The proteogenomic subtypes of acute myeloid leukemia. Cancer Cell 40, 301–317.e12 (2022).

30. Salovska, B. et al. Peroxiredoxin 6 protects irradiated cells from oxidative stress and shapes their senescence-associated cytokine landscape. Redox Biol. 49, 102212 (2022).

31. Gunter, H. M. et al. A universal molecular control for DNA, mRNA and protein expression. Nat. Commun. 15, 2480 (2024).

32. Bradley, D. et al. The fitness cost of spurious phosphorylation. EMBO J. 43, 4720–4751 (2024).

33. Guzman, U. H. et al. Ultra-fast label-free quantification and comprehensive proteome coverage with narrow-window data-independent acquisition. Nat. Biotechnol. 42, 1855–1866 (2024).

34. Kattelus, R. et al. Phenotypic profiling of human induced regulatory T cells at early differentiation: insights into distinct immunosuppressive potential. Cell. Mol. Life Sci. 81, 399 (2024).

35. Zare, A. et al. Axonal tau reduction ameliorates tau and amyloid pathology in a mouse model of Alzheimer’s disease. Transl. Neurodegener. 14, 39 (2025).

36. Zhang, H. et al. Heterochromatome wide analyses reveal MBD2 as a phase separation scaffold for heterochromatin compartmentalization and composition. Nucleic Acids Res. 53, gkaf1380 (2025).

37. Romero-Pérez, P. S. et al. Protein surface chemistry encodes an adaptive tolerance to desiccation. Cell Syst. 16, 101407 (2025).

38. Arroyo-Gomez, J. et al. Functional landscape of ubiquitin linkages couples K29-linked ubiquitylation to epigenome integrity. EMBO J. 44, 6944–6978 (2025).

39. Botella, J. et al. Sprint interval exercise disrupts mitochondrial ultrastructure driving a unique mitochondrial stress response and remodelling in men. Nat. Commun. 17, 71 (2025).

40. Pereyra, G. et al. SFRP1 upregulation causes hippocampal synaptic dysfunction and memory impairment. Cell Rep. 44, 115535 (2025).

41. Gharibi, B. et al. Post-gastrulation amnioids as a stem cell-derived model of human extra-embryonic development. Cell 188, 3757–3774.e20 (2025).

42. Su, J. et al. Polymerization-mediated SRFR1 condensation in upper lateral root cap cells regulates root growth. Plant Cell 38, koaf292 (2026).

43. Benjamini, Y. & Hochberg, Y. Controlling the False Discovery Rate: A Practical and Powerful Approach to Multiple Testing. J. R. Stat. Soc. Ser. B Methodol. 57, 289–300 (1995).

44. Hinterholzer, A. et al. Detecting aspartate isomerization and backbone cleavage after aspartate in intact proteins by NMR spectroscopy. J. Biomol. NMR 75, 71–82 (2021).

45. Bodenmiller, B. et al. Phosphoproteomic Analysis Reveals Interconnected System-Wide Responses to Perturbations of Kinases and Phosphatases in Yeast. Sci. Signal. 3, rs4–rs4 (2010).

46. Bekker-Jensen, D. B. et al. Rapid and site-specific deep phosphoproteome profiling by data-independent acquisition without the need for spectral libraries. Nat. Commun. 11, 787 (2020).

47. Wang, Y. et al. GABAA receptor π forms channels that stimulate ERK through a G-protein-dependent pathway. Mol. Cell 85, 166–176.e5 (2025).

48. He, Y. et al. Evaluation of the Orbitrap Ascend Tribrid Mass Spectrometer for Shotgun Proteomics. Anal. Chem. 95, 10655–10663 (2023).

49. Criscuolo, A., Zeller, M. & Fedorova, M. Evaluation of Lipid In-Source Fragmentation on Different Orbitrap-based Mass Spectrometers. J. Am. Soc. Mass Spectrom. 31, 463–466 (2020).

50. Yu, F. et al. Fast Quantitative Analysis of timsTOF PASEF Data with MSFragger and IonQuant. Mol. Cell. Proteomics 19, 1575–1585 (2020).

51. Reber, V. et al. Paradoxical non-catalytic kinase functions are driven by inhibitor-induced displacement of autoinhibitory domains. 2025.11.06.687012 Preprint at 10.1101/2025.11.06.687012 (2026).

52. Elsässer, F. et al. Limited proteolysis-coupled mass spectrometry captures proteome-wide protein structural alterations and biomolecular condensation in living cells. Mol. Syst. Biol. (2026) doi:10.1038/s44320-025-00182-6.

53. Abelin, J. G. & Cox, A. L. Innovations Toward Immunopeptidomics. Mol. Cell. Proteomics 23, 100823 (2024).

54. Purcell, A. W., Ramarathinam, S. H. & Ternette, N. Mass spectrometry–based identification of MHC-bound peptides for immunopeptidomics. Nat. Protoc. 14, 1687–1707 (2019).

55. Pak, H. et al. Sensitive Immunopeptidomics by Leveraging Available Large-Scale Multi-HLA Spectral Libraries, Data-Independent Acquisition, and MS/MS Prediction. Mol. Cell. Proteomics 20, 100080 (2021).

56. Kessler, A. L. et al. HLA I immunopeptidome of synthetic long peptide pulsed human dendritic cells for therapeutic vaccine design. Npj Vaccines 10, 12 (2025).

57. Dorvash, M., Illing, P. T., Croft, N. P., Ramarathinam, S. H. & Purcell, A. W. Deep Exploration of the Immunopeptidome of a Pancreatic Cancer Cell Line: Implications for Clinical Immunopeptidomics and Immunotherapy. Mol. Cell. Proteomics 24, 101030 (2025).

58. Tanuwidjaya, E. et al. SAPrIm 2.0: a semi-automated protocol for mid-throughput soluble HLA immunopeptidomics. Front. Immunol. 16, (2025).

59. Cappelletti, V. et al. Dynamic 3D proteomes reveal protein functional alterations at high resolution in situ. Cell 184, 545–559.e22 (2021).

60. Mehta, V. et al. Structure of Mycobacterium tuberculosis Cya, an evolutionary ancestor of the mammalian membrane adenylyl cyclases. eLife 11, e77032 (2022).

61. Li, K. et al. A peptide-centric local stability assay enables proteome-scale identification of the protein targets and binding regions of diverse ligands. Nat. Methods 22, 278–282 (2025).

62. Schubert, O. T. et al. Building high-quality assay libraries for targeted analysis of SWATH MS data. Nat. Protoc. 10, 426–441 (2015).

63. Dunham, W. H., Mullin, M. & Gingras, A.-C. Affinity-purification coupled to mass spectrometry: Basic principles and strategies. PROTEOMICS 12, 1576–1590 (2012).

64. O’Reilly, F. J. & Rappsilber, J. Cross-linking mass spectrometry: methods and applications in structural, molecular and systems biology. Nat. Struct. Mol. Biol. 25, 1000–1008 (2018).

65. Goel, R. K., Bithi, N. & Emili, A. Trends in co-fractionation mass spectrometry: A new gold-standard in global protein interaction network discovery. Curr. Opin. Struct. Biol. 88, 102880 (2024).

66. Malta, C. F. et al. Pushing the limits of hydrogen/deuterium exchange mass spectrometry to study protein:fragment low affinity interactions. Commun. Chem. 8, 405 (2025).

67. Harvey, D. J. Analysis of Protein Glycosylation by Mass Spectrometry. in Analysis of Protein Post-Translational Modifications by Mass Spectrometry 89–159 (John Wiley & Sons, Ltd, 2016). doi:10.1002/9781119250906.ch3.

68. Cardon, T., Fournier, I. & Salzet, M. Chasing the Ghost Proteome in the Dark Matter. Mol. Cell. Proteomics 24, 101076 (2025).

69. Varjosalo, M. et al. The Protein Interaction Landscape of the Human CMGC Kinase Group. Cell Rep. 3, 1306–1320 (2013).

70. The UniProt Consortium. UniProt: the Universal Protein Knowledgebase in 2025. Nucleic Acids Res. 53, D609–D617 (2025).

71. Gamble, D. venn (https://ch.mathworks.com/matlabcentral/fileexchange/22282-venn), MATLAB Central File Exchange. (2026).

