## Supplementary Information for "In-source fragmentation in mass spectrometry-based proteomics: prevalence, impact, and strategies for mitigation"

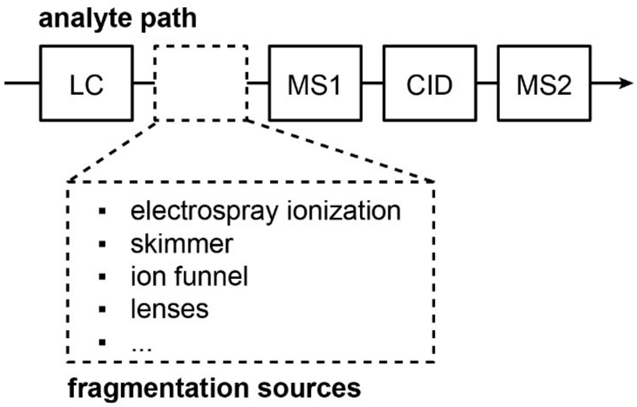


**Supplementary Fig. 1. Conceptual figure on the location of ISF in the analyte path during MS.** ISF occurs after liquid chromatography (LC) and prior to the first mass filter (MS1) during LC-MS. Sources of ISF can be electrospray ionization but also electric fields for ion routing like in the ion funnel. CID: collision induced dissociation

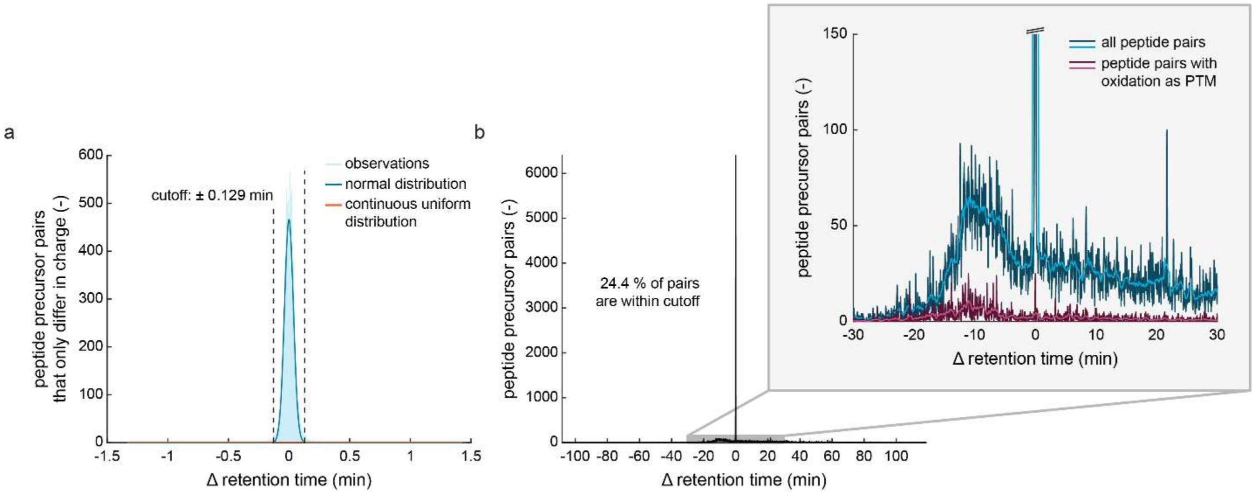


**Supplementary Fig. 2. Examples of ΔRT distributions for cutoff estimation and application.**

**a,** Histogram showing the number of peptide pairs that only differ in charge per ΔRT (min) (number of bins = 1000). Light blue shade indicates measured data from a yeast reference sample spiked with 8 pure proteins (n = 1 because the ISF detection relies on individual samples by design). Dark blue line indicates a normal distribution fitted to the data, red a continuous uniform distribution. The cutoff (dotted lines) is defined as the ΔRT at which both model distributions intersect.

**b,** Histograms showing the number of peptide pairs that do not differ in charge per ΔRT (min) in the sample of a (number of bins = 5000). Graph in the grey box shows an excerpt of the plot outside. Data shown in blue indicates all peptide pairs (except peptides only differing in charge), data in red all peptide pairs that feature an oxidation on methionine as PTM. The light blue and red lines indicate smoothing (moving average with a window size = 20) over the data (dark blue and red, respectively).

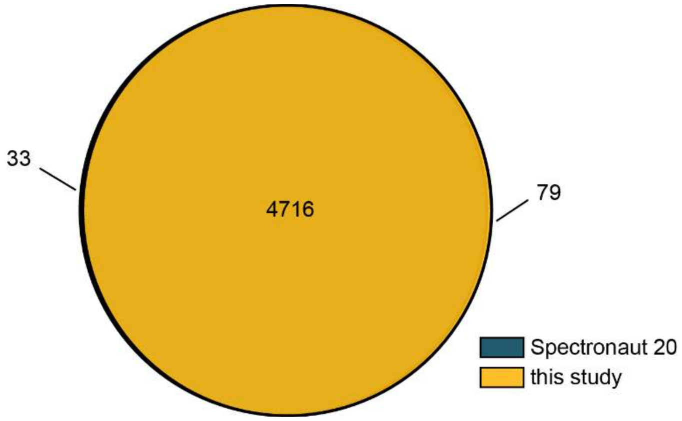


**Supplementary Fig. 3. Comparison of ISF detection algorithms.**
Venn diagram showing the number of ISF annotations across n = 4 technical replicates of our yeast reference sample spiked with 8 pure proteins. The yellow circle indicates the ISF annotations by the ISF detection algorithm presented in this study. The blue circle indicates the ISF annotations by the ISF detection algorithm by Spectronaut 20 (Biognosys, Schlieren, Switzerland).

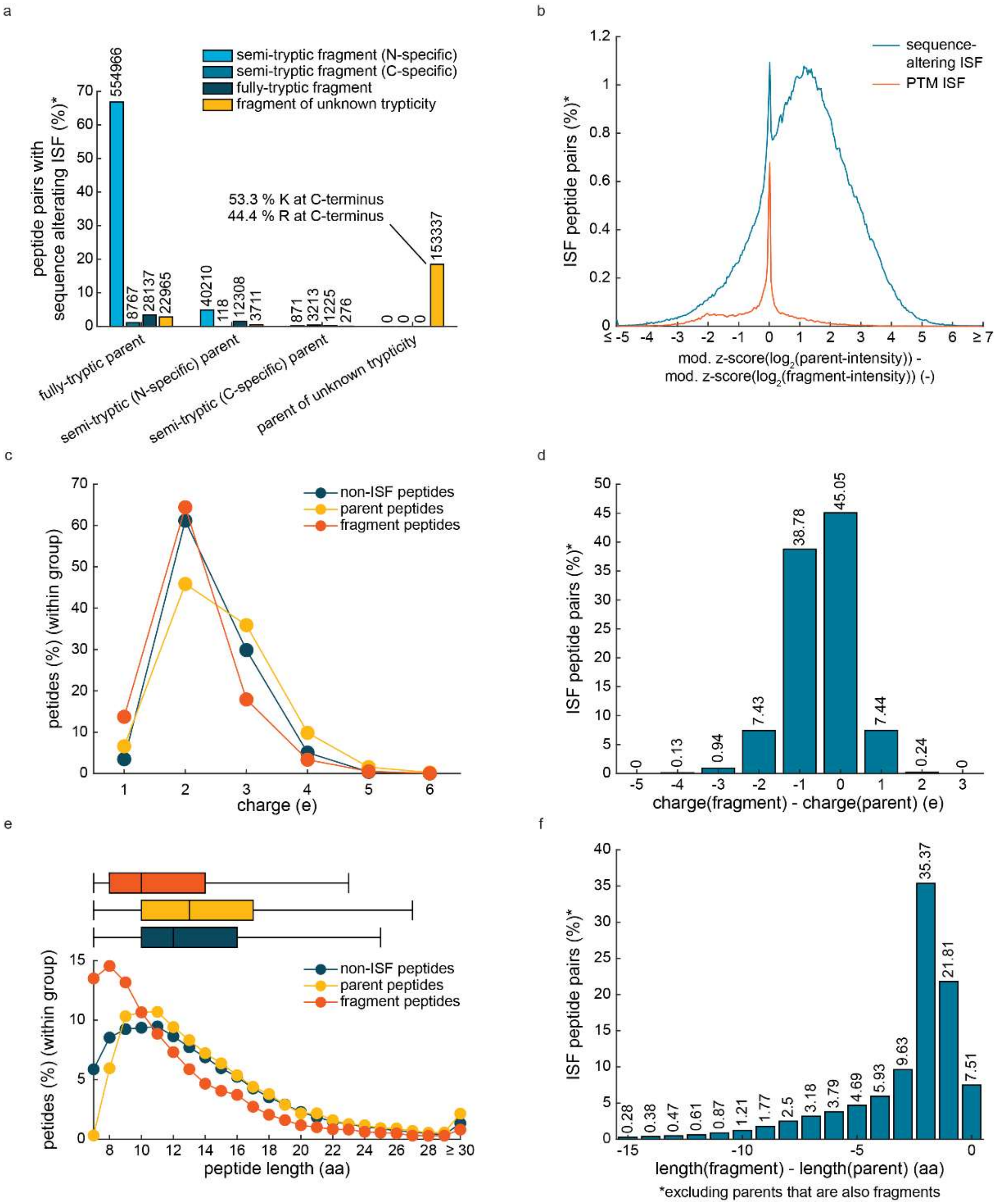


**Supplementary Fig. 4. Properties of in-source fragments**

**a,** Grouped bar plot showing the number of peptide pairs with sequencing-altering ISF (*excluding fragment-to-fragment pairs). Each group of bars indicate the trypticity of the ISF parent, and bars within groups the apparent trypticity of the respective in-source fragments. For the entire figure, the data stems from 23 combined datasets (also see Fig. 3). In-source fragments are not due to trypsin semi- or unspecific cleavages however they appear as such peptides. Light blue bars indicate trypsin N-specific in-source fragments (blue = C-specific in-source fragments, dark blue = fully tryptic in-source fragments). Yellow bars indicate peptide pairs for which trypticity could be unambiguously determined as the parent peptide was non-proteotypic.

**b,** Histogram showing the number of peptide pairs with sequence-altering ISF (%, blue line) and PTM ISF (%, red line) per difference in z-scored (modified) $log_2$-intensities between ISF parents and fragments (number of bins = 300). The data does not contain pairs between two in-source fragments. Modified z-scores rely on median values and median absolute deviations and thus are more robust against outliers than conventional z-scores.

**c,** Dot plot showing the number of peptides (%, normalized per group) per charge. Data in blue indicates peptides without evidence of ISF (yellow = ISF parents, red = in-source fragments).

**d,** Bar plot showing the number of peptide pairs with evidence of ISF (%) (excluding pairs between two in-source fragments) per difference in charge between in-source fragments and parents.

**e,** Dot plot showing the number of peptides (% per group) per peptide length. Data in blue indicates peptides without evidence of ISF (yellow =ISF parents, red = in-source fragments). Box whisker plots above the distribution show the median values, boxes depict interquartile ranges, and whiskers extend to 1.5-fold of the interquartile ranges.

**f,** Bar plot showing the number of peptide pairs with evidence of ISF (%) (excluding pairs between two in-source fragments) per difference in peptide length between in-source fragments and parents.

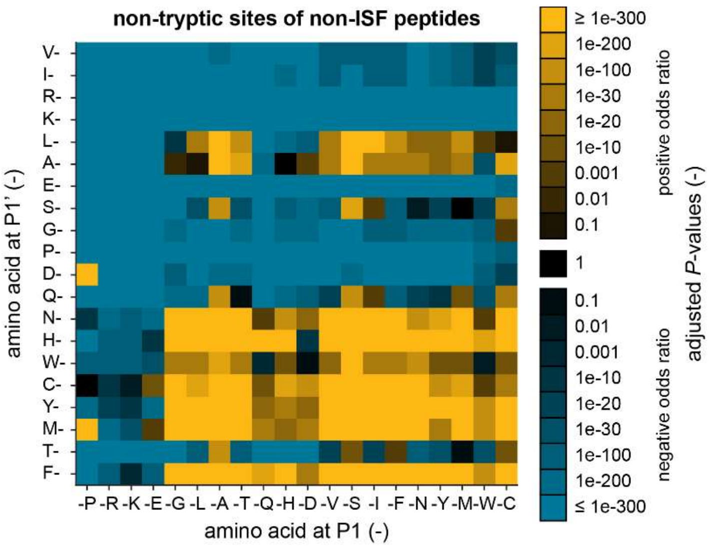


**Supplementary Fig. 5. Cleavage site enrichment analysis of non-ISF peptides**

Using the combined 23 datasets shown in Fig. 2, we analyzed all non-tryptic cleavage sites (N- and C-terminal) of non-ISF peptides. P1 denotes the amino acid C-terminal of a cleavage site, and P1' the amino acid prior to the cleavage site. The heatmap shows adjusted *P*-values from an enrichment analysis (Fisher's exact test, two-tailed). *P*-values were adjusted using the procedure by Benjamini-Hochberg.[1] Data marked in yellow indicates positive odds ratios, blue negative odds ratios.

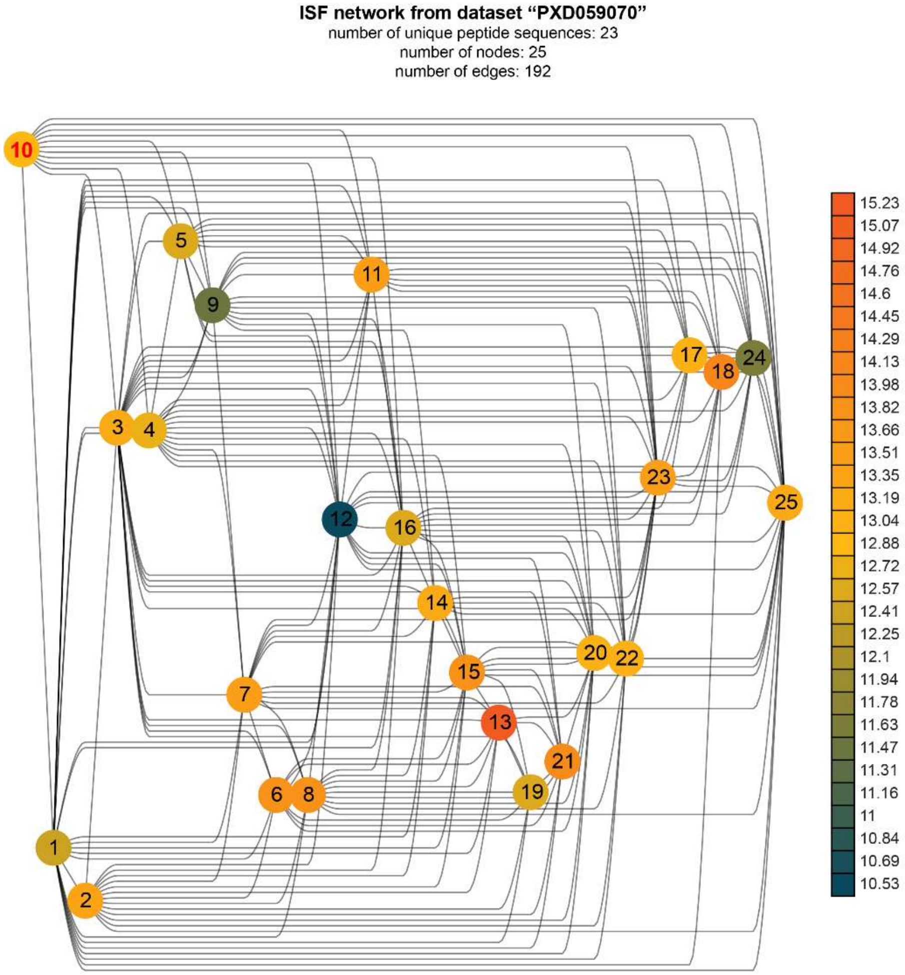


| | |
|---|---|
| 1 | GGGHSGGSGGGHSGGSGGNYGGGSGSGGGSGGGYGGGSGSR.4 |
| 2 | GGGHSGGSGGNYGGGSGSGGGSGGGYGGGSGSR.3 |
| 3 | GGGSGGGHSGGSGGGHSGGSGGNYGGGSGSGGGSGGGYGGGSGSR.4 |
| 4 | GGGYGGGSGGGHSGGSGGGHSGGSGGNYGGGSGSGGGSGGGYGGGSGSR.4 |
| 5 | GGHSGGSGGGHSGGSGGNYGGGSGSGGGSGGGYGGGSGSR.3 |
| 6 | GGHSGGSGGGHSGGSGGNYGGGSGSGGGSGGGYGGGSGSR.4 |
| 7 | GGHSGGSGGNYGGGSGSGGGSGGGYGGGSGSR.3 |
| 8 | GGSGGGHSGGSGGGHSGGSGGNYGGGSGSGGGSGGGYGGGSGSR.4 |
| 9 | GGSGGGHSGGSGGNYGGGSGSGGGSGGGYGGGSGSR.3 |
| **10** | **GGSGGGYGGGSGGGHSGGSGGGHSGGSGGNYGGGSGSGGGSGGGYGGGSGSR.4** |
| 11 | GGYGGGSGGGHSGGSGGGHSGGSGGNYGGGSGSGGGSGGGYGGGSGSR.4 |
| 12 | GHSGGSGGGHSGGSGGNYGGGSGSGGGSGGGYGGGSGSR.3 |
| 13 | GHSGGSGGGHSGGSGGNYGGGSGSGGGSGGGYGGGSGSR.4 |
| 14 | GHSGGSGGNYGGGSGSGGGSGGGYGGGSGSR.3 |
| 15 | GSGGGHSGGSGGGHSGGSGGNYGGGSGSGGGSGGGYGGGSGSR.4 |
| 16 | GSGGGHSGGSGGNYGGGSGSGGGSGGGYGGGSGSR.3 |
| 17 | GSGGGYGGGSGGGHSGGSGGGHSGGSGGNYGGGSGSGGGSGGGYGGGSGSR.4 |
| 18 | GYGGGSGGGHSGGSGGGHSGGSGGNYGGGSGSGGGSGGGYGGGSGSR.4 |
| 19 | HSGGSGGGHSGGSGGNYGGGSGSGGGSGGGYGGGSGSR.4 |
| 20 | HSGGSGGNYGGGSGSGGGSGGGYGGGSGSR.3 |
| 21 | SGGGHSGGSGGGHSGGSGGNYGGGSGSGGGSGGGYGGGSGSR.4 |
| 22 | SGGGHSGGSGGNYGGGSGSGGGSGGGYGGGSGSR.3 |
| 23 | SGGGYGGGSGGGHSGGSGGGHSGGSGGNYGGGSGSGGGSGGGYGGGSGSR.4 |
| 24 | SGGNYGGGSGSGGGSGGGYGGGSGSR.2 |
| 25 | SGGSGGGHSGGSGGNYGGGSGSGGGSGGGYGGGSGSR.3 |

**Supplementary Fig. 6. Example of an ISF network.**

Network graph showing the largest ISF network detected in the combined 23 datasets from Fig. 2. Nodes indicate peptide precursors, edges connected peptide precursors with evidence of ISF. The color of nodes indicates the $log_2$-mean intensity of the respective peptide precursors (n = 45 samples). The peptide precursor highlighted by bold, red font is the longest peptide and thus, by definition, the only ISF parent. Numbers after peptide precursor sequences indicate the number of charges.

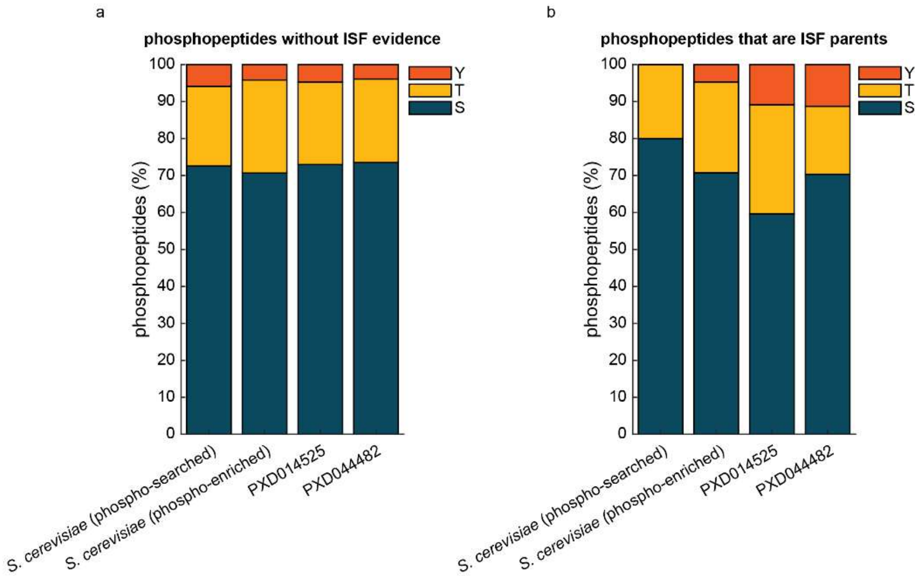


**Supplementary Fig. 7. The number of phosphopeptides by their modified amino acid.**

**a,** Bar graph showing the number of phosphopeptides without evidence of ISF (%) with a phosphorylated tyrosine (indicated by red bars), threonine (yellow bars), and serine (blue bars) in four different datasets as indicated.

**b,** same as A but, instead of data from phosphopeptides without evidence of ISF, data of phosphopeptides that are ISF parents is shown.

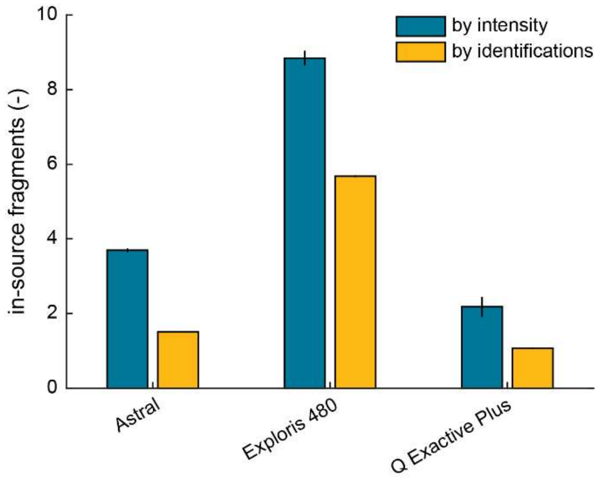


**Supplementary Fig. 8. Comparison of the amount of ISF between different mass spectrometers.** Bar plot showing the average percentage of in-source fragments (n = 4) in data from the same sample acquired on three different Thermo Scientific mass spectrometers: Orbitrap Exploris 480, Orbitrap Astral, Q Exactive Plus Hybrid Quadrupole-Orbitrap (QE+). Yellow bars indicate number of identifications (%), blue bars intensities (%). Vertical black lines indicate the standard deviation.

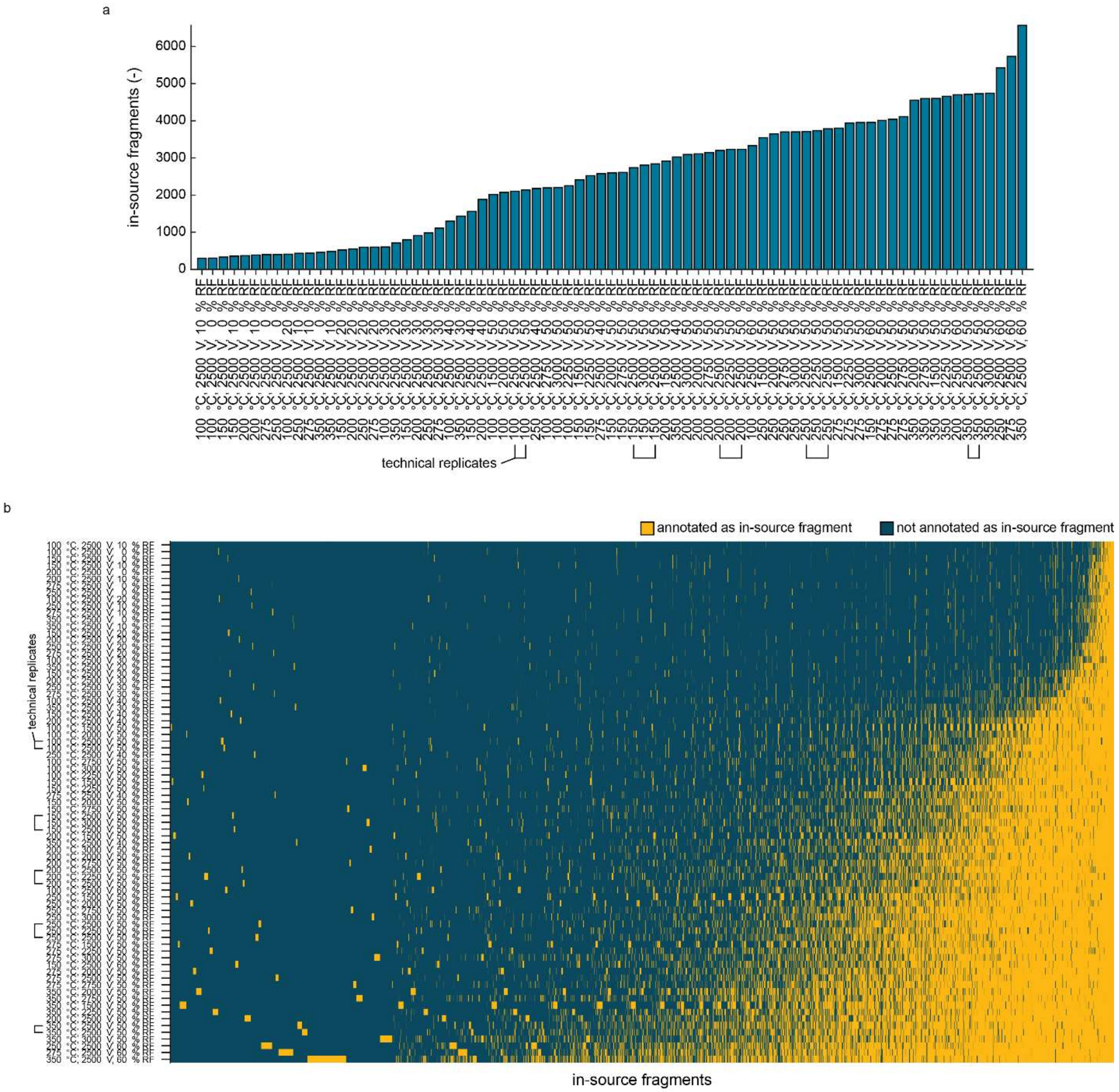


**Supplementary Fig. 9. Effect of MS parameters on ISF.**

**a,** Bar graph showing the number of identified in-source fragments in a yeast reference sample spiked with 8 pure proteins acquired with different electrospray voltages, ITC temperatures, and funnel radiofrequencies (n = 1, except 5 pairs of technical duplicates as indicated).

**b,** Heatmap showing whether a peptide, which has been identified at least once as an in-source fragment across all samples shown in a, is identified as an in-source fragment in each of the 77 samples (yellow = peptide is identified as in-source fragment, blue = peptide is not identified as an in-source fragment). Samples are sorted from low amount of ISF (top) to high (bottom). Peptides are sorted from a low number of identifications as an in-source fragment across all samples (left) to high (right).

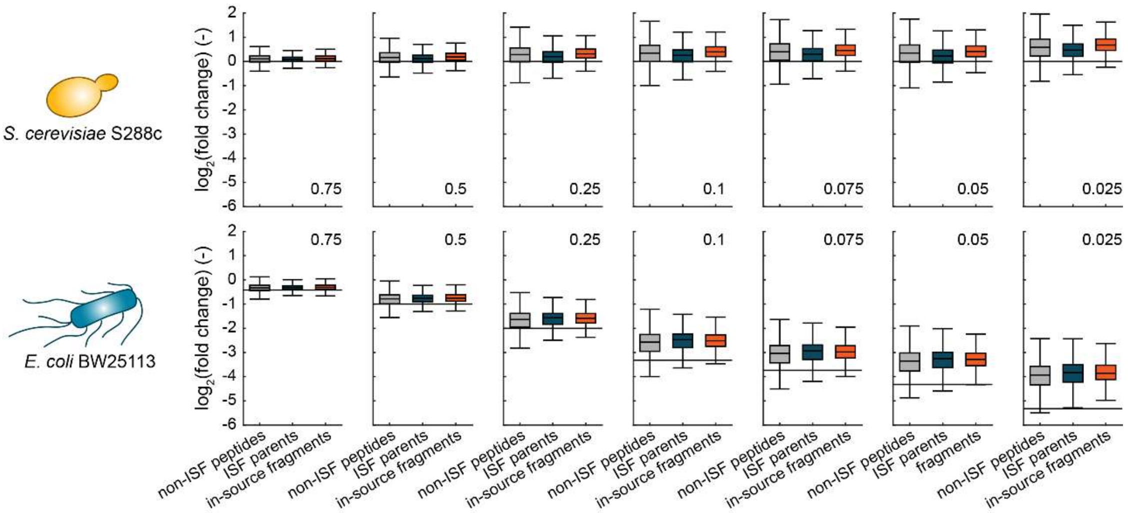


**Supplementary Fig. 10.**

Box whisker plot depicting median values of the data shown in Fig. 6.b. Boxes represent interquartile ranges, and whiskers extend to 1.5-fold of the interquartile ranges. The numbers within the plots indicate the respective dilution factors of the *E. coli* proteome (also see Fig. 6.a). Grey boxes indicate data from non-ISF peptides (blue boxes = ISF parents, red boxes = in-source fragments). Horizontal black lines show the theoretical $log_2$-fold changes that are expected due to the experimental design. The upper panel of figures displays the data of the yeast peptides, the lower panel the data of the *E. coli* peptides.

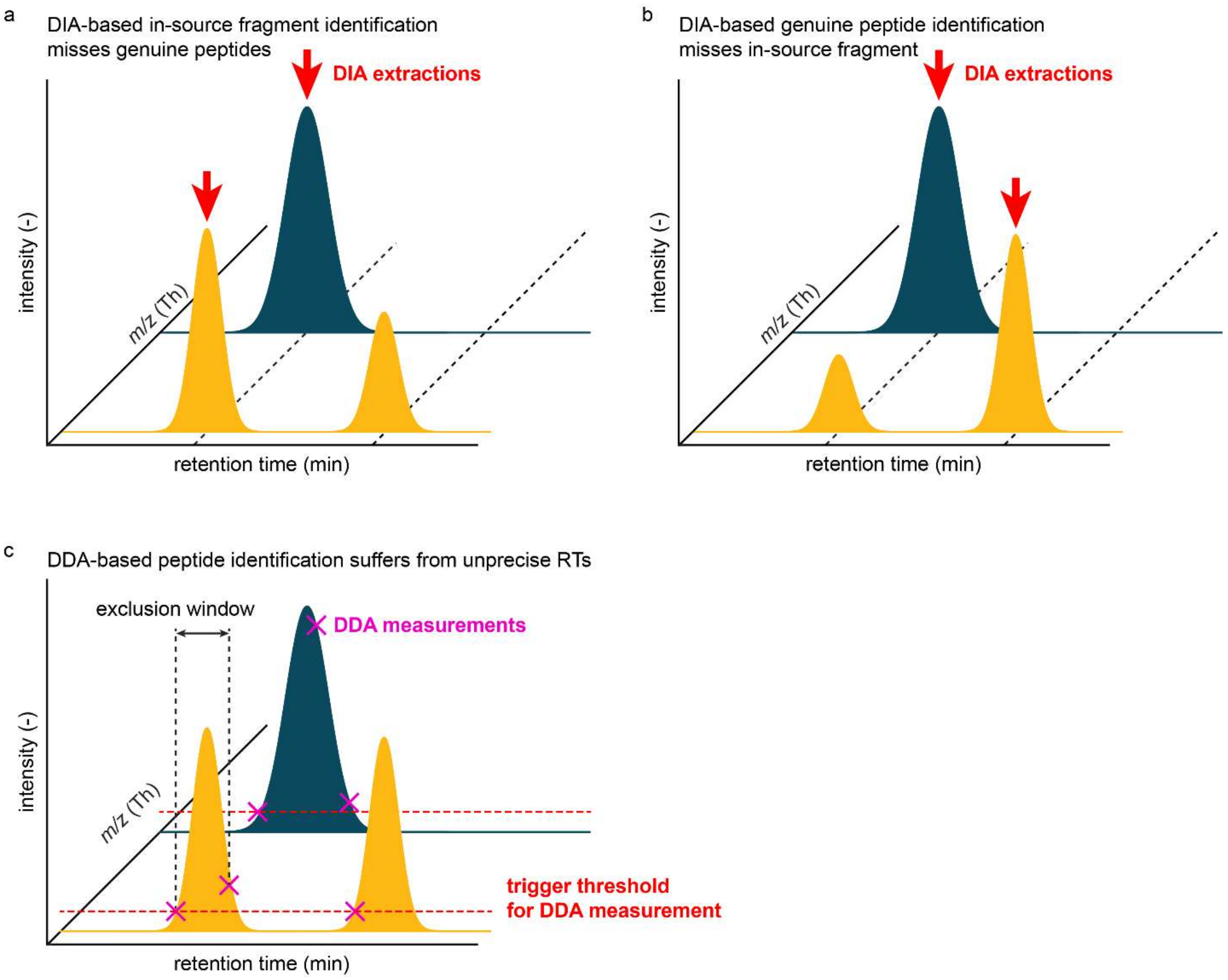


**Supplementary Fig. 11. The implications of the one-peak assumption of current DIA extraction approaches on data with ISF.**

**a,** Conceptual figure showing hypothetical elution profiles of peptides from DIA data. The x-axis shows the mass over charge *m/z* (Th), the y-axis the retention time (min), and the z-axis the intensity (-). Some elution profiles feature two peaks, like the yellow profile in our example, or even more. Out of such peaks, some will be due to ISF of a parent peptide (indicated by the blue elution profile). During DIA extraction, current search tools allow only a single peak for extraction, which can lead to missing the identification of genuine peptides if the peak of an in-source fragment is chosen (peak selection is indicated by red arrows).

**b,** Like a, but instead of the in-source fragment, the genuine peptide is selected for DIA extraction, which leads to an underestimation of ISF.

**c,** Like a, but instead of DIA data, DDA data is considered. If the signal of a peak reaches a certain threshold, a MS2 measurement will be triggered. Depending on the peak width and the size of the exclusion window, peaks can be covered inconsistently making precise detection of peak apices and, subsequently, ISF difficult.

**Supplementary Tab. 1. Literature datasets used in this study.**

| PRIDE ID | Author | DOI | Mass spectrometer | Sample source | Type |
|---|---|---|---|---|---|
| **PXD014525** | Bekker-Jensen *et al.*[2] | https://doi.org/10.1038/s41467-020-14609-1 | Thermo Fisher Scientific Q Exactive HF-X | *H. sapiens* | phosphoproteomics |
| **PXD019777** | Kalxdorf *et al.*[3] | https://doi.org/10.1038/s41467-021-25077-6 | Thermo Fisher Scientific Q Exactive HF | *E. coli* | trypsin-digested bottom-up proteomics |
| **PXD020761** | Midha *et al.*[4] | https://doi.org/10.1038/s41597-020-00724-7 | SCIEX TripleTOF 6600 | *E. coli* | trypsin-digested bottom-up proteomics |
| **PXD022297** | Cappelletti *et al.*[5] | https://doi.org/10.1016/j.cell.2020.12.021 | Thermo Fisher Scientific Q Exactive Plus | *S. cerevisiae* | LiP dataset |
| **PXD022950** | Pak *et al.*[6] | https://doi.org/10.1016/j.mcpro.2021.100080 | Thermo Fisher Scientific Q Exactive HF-X | *H. sapiens* | immunopeptidomics |
| **PXD028007** | Jayavelu *et al.*[7] | https://doi.org/10.1016/j.ccell.2022.02.006 | Thermo Fisher Scientific Q Exactive HF-X | *H. sapiens* | trypsin-digested bottom-up proteomics |
| **PXD028772** | Salovska *et al.*[8] | https://doi.org/10.1016/j.redox.2021.102212 | Thermo Fisher Scientific Exploris 480 | *H. sapiens* | trypsin-digested bottom-up proteomics |
| **PXD033826** | Mehta *et al.*[9] | https://doi.org/10.7554/eLife.77032 | Thermo Fisher Scientific Exploris 480 | *E. coli* | LiP dataset |
| **PXD034606** | Li *et al.*[10] | https://doi.org/10.1038/s41592-024-02553-7 | Thermo Fisher Scientific Exploris 480 | *H. sapiens* | LiP dataset |
| **PXD035035** | Gunter *et al.*[11] | https://doi.org/10.1038/s41467-024-46456-9 | Thermo Fisher Scientific Q Exactive HF | *E. coli* | trypsin-digested bottom-up proteomics |
| **PXD044482** | Wang *et al.*[12] | https://doi.org/10.1016/j.molcel.2024.11.016 | Thermo Fisher Scientific Fusion Lumos Tribrid | *H. sapiens* | phosphoproteomics |
| **PXD045466** | Bradley *et al.*[13] | https://doi.org/10.1038/s44318-024-00200-7 | Thermo Fisher Scientific Exploris 480 | *S. cerevisiae* | trypsin-digested bottom-up proteomics |
| **PXD046444** | Guzman *et al.*[14] | https://doi.org/10.1038/s41587-023-02099-7 | Thermo Fisher Scientific Astral | *H. sapiens, S. cerevisiae, E. coli* | trypsin-digested bottom-up proteomics |
| **PXD051490** | Kessler *et al.*[15] | https://doi.org/10.1038/s41541-025-01069-1 | Thermo Fisher Scientific Eclipse Tribrid | *H. sapiens* | immunopeptidomics |
| **PXD052626** | Kattelus *et al.* | https://doi.org/10.1007/s00018-024-05429-3 | Thermo Fisher Scientific Fusion Lumos Tribrid with FAIMS | *H. sapiens* | trypsin-digested bottom-up proteomics |
| **PXD053387** | Zare *et al.*[16] | https://doi.org/10.1186/s40035-025-00499-0 | Thermo Fisher Scientific Astral with FAIMS | *M. muculus* | trypsin-digested bottom-up proteomics |
| **PXD054417** | Dorvash *et al.* | https://doi.org/10.1016/j.mcpro.2025.101030 | SCIEX 7600 ZenoTOF | *H. sapiens* | immunopeptidomics |
| **PXD055626** | Zhang *et al.*[17] | https://doi.org/10.1093/nar/gkaf1380 | Thermo Fisher Scientific Exploris 480 | *M. muculus* | trypsin-digested bottom-up proteomics |
| **PXD055855** | Romero-Pérez *et al.*[18] | https://doi.org/10.1016/j.cels.2025.101407 | Bruker timsTOF Pro2 | *S. cerevisiae* | trypsin-digested bottom-up proteomics |
| **PXD056564** | Arroyo-Gomez *et al.*[19] | https://doi.org/10.1038/s44318-025-00599-7 | Thermo Fisher Scientific Astral | *H. sapiens* | trypsin-digested bottom-up proteomics |
| **PXD058880** | Tanuwidjaya *et al.*[20] | https://doi.org/10.3389/fimmu.2025.1546629 | Thermo Fisher Scientific Exploris 480 | *H. sapiens* | immunopeptidomics |
| **PXD059070** | Botella *et al.*[21] | https://doi.org/10.1038/s41467-025-66625-8 | Thermo Fisher Scientific Eclipse Tribrid | *H. sapiens* | trypsin-digested bottom-up proteomics |
| **PXD060276** | Pereyra *et al.*[22] | https://doi.org/10.1016/j.celrep.2025.115535 | Bruker timsTOF Pro2 | *M. muculus* | trypsin-digested bottom-up proteomics |
| **PXD062917** | Gharibi *et al.*[23] | https://doi.org/10.1016/j.cell.2025.04.025 | Bruker timsTOF Pro2 | *H. sapiens* | trypsin-digested bottom-up proteomics |
| **PXD069457** | Su *et al.*[24] | https://doi.org/10.1093/plcell/koaf292 | Bruker timsTOF Pro2 | *A. thaliana* | trypsin-digested bottom-up proteomics |

### References


1. Benjamini, Y. & Hochberg, Y. Controlling the False Discovery Rate: A Practical and Powerful Approach to Multiple Testing. *J. R. Stat. Soc. Ser. B Methodol.* **57**, 289–300 (1995).
2. Bekker-Jensen, D. B. *et al.* Rapid and site-specific deep phosphoproteome profiling by data-independent acquisition without the need for spectral libraries. *Nat. Commun.* **11**, 787 (2020).
3. Kalxdorf, M., Müller, T., Stegle, O. & Krijgsveld, J. IceR improves proteome coverage and data completeness in global and single-cell proteomics. *Nat. Commun.* **12**, 4787 (2021).
4. Midha, M. K. *et al.* A comprehensive spectral assay library to quantify the Escherichia coli proteome by DIA/SWATH-MS. *Sci. Data* **7**, 389 (2020).
5. Cappelletti, V. *et al.* Dynamic 3D proteomes reveal protein functional alterations at high resolution in situ. *Cell* **184**, 545-559.e22 (2021).
6. Pak, H. *et al.* Sensitive Immunopeptidomics by Leveraging Available Large-Scale Multi-HLA Spectral Libraries, Data-Independent Acquisition, and MS/MS Prediction. *Mol. Cell. Proteomics* **20**, 100080 (2021).
7. Jayavelu, A. K. *et al.* The proteogenomic subtypes of acute myeloid leukemia. *Cancer Cell* **40**, 301-317.e12 (2022).
8. Salovska, B. *et al.* Peroxiredoxin 6 protects irradiated cells from oxidative stress and shapes their senescence-associated cytokine landscape. *Redox Biol.* **49**, 102212 (2022).
9. Mehta, V. *et al.* Structure of Mycobacterium tuberculosis Cya, an evolutionary ancestor of the mammalian membrane adenylyl cyclases. *eLife* **11**, e77032 (2022).
10. Li, K. *et al.* A peptide-centric local stability assay enables proteome-scale identification of the protein targets and binding regions of diverse ligands. *Nat. Methods* **22**, 278–282 (2025).
11. Gunter, H. M. *et al.* A universal molecular control for DNA, mRNA and protein expression. *Nat. Commun.* **15**, 2480 (2024).

12. Wang, Y. *et al.* GABAA receptor π forms channels that stimulate ERK through a G-protein-dependent pathway. *Mol. Cell* **85**, 166-176.e5 (2025).
13. Bradley, D. *et al.* The fitness cost of spurious phosphorylation. *EMBO J.* **43**, 4720–4751 (2024).
14. Guzman, U. H. *et al.* Ultra-fast label-free quantification and comprehensive proteome coverage with narrow-window data-independent acquisition. *Nat. Biotechnol.* **42**, 1855–1866 (2024).
15. Kessler, A. L. *et al.* HLA I immunopeptidome of synthetic long peptide pulsed human dendritic cells for therapeutic vaccine design. *Npj Vaccines* **10**, 12 (2025).
16. Zare, A. *et al.* Axonal tau reduction ameliorates tau and amyloid pathology in a mouse model of Alzheimer's disease. *Transl. Neurodegener.* **14**, 39 (2025).
17. Zhang, H. *et al.* Heterochromatome wide analyses reveal MBD2 as a phase separation scaffold for heterochromatin compartmentalization and composition. *Nucleic Acids Res.* **53**, gkaf1380 (2025).
18. Romero-Pérez, P. S. *et al.* Protein surface chemistry encodes an adaptive tolerance to desiccation. *Cell Syst.* **16**, 101407 (2025).
19. Arroyo-Gomez, J. *et al.* Functional landscape of ubiquitin linkages couples K29-linked ubiquitylation to epigenome integrity. *EMBO J.* **44**, 6944–6978 (2025).
20. Tanuwidjaya, E. *et al.* SAPrIm 2.0: a semi-automated protocol for mid-throughput soluble HLA immunopeptidomics. *Front. Immunol.* **16**, (2025).
21. Botella, J. *et al.* Sprint interval exercise disrupts mitochondrial ultrastructure driving a unique mitochondrial stress response and remodelling in men. *Nat. Commun.* **17**, 71 (2025).
22. Pereyra, G. *et al.* SFRP1 upregulation causes hippocampal synaptic dysfunction and memory impairment. *Cell Rep.* **44**, 115535 (2025).
23. Gharibi, B. *et al.* Post-gastrulation amnioids as a stem cell-derived model of human extra-embryonic development. *Cell* **188**, 3757-3774.e20 (2025).

24. Su, J. *et al.* Polymerization-mediated SRFR1 condensation in upper lateral root cap cells regulates root growth. *Plant Cell* **38**, koaf292 (2026).
